# Reduced Glucose Sensation Can Increase the Fitness of *Saccharomyces cerevisiae* Lacking Mitochondrial DNA

**DOI:** 10.1101/024331

**Authors:** Emel Akdoğan, Mehmet Tardu, Görkem Garipler, Gülkız Baytek, İ. Halil Kavaklı, Cory D. Dunn

## Abstract

Damage to the mitochondrial genome (mtDNA) can lead to diseases for which there are no clearly effective treatments. Since mitochondrial function and biogenesis are controlled by the nutrient environment of the cell, it is possible that perturbation of conserved, nutrient-sensing pathways may successfully treat mitochondrial disease. We found that restricting glucose or otherwise reducing the activity of the protein kinase A (PKA) pathway can lead to improved proliferation of *Saccharomyces cerevisiae* cells lacking mtDNA and that the transcriptional response to mtDNA loss is reduced in cells with diminished PKA activity. We have excluded many pathways and proteins from being individually responsible for the benefits provided to cells lacking mtDNA by PKA inhibition, and we found that robust import of mitochondrial polytopic membrane proteins may be required in order for cells without mtDNA to receive the full benefits of PKA reduction. Finally, we have discovered that the transcription of genes involved in arginine biosynthesis and aromatic amino acid catabolism is altered after mtDNA damage. Our results highlight the potential importance of nutrient detection and availability on the outcome of mitochondrial dysfunction.

## Introduction

Mitochondria are the location of ATP synthesis by oxidative phosphorylation (OXPHOS). In addition, essential biosynthetic pathways, such as iron-sulfur cluster biogenesis [1,2], are compartmentalized within mitochondria. Genetic material retained from a bacterial ancestor [3] supports the process of OXPHOS. Proteins required to generate a proton gradient across the mitochondrial inner membrane (IM) are encoded by mitochondrial DNA (mtDNA), as are proteins allowing this proton gradient to power ATP synthesis [4]. In humans, pathological mutations of mtDNA can be inherited [5] or may accumulate following pharmacological treatment for viral infections [6] or cancer [7,8]. Many organisms, including humans, accumulate cells containing significant levels of damaged mtDNA during their lifespan, and it is therefore possible that mtDNA mutations can promote the aging process [9,10].

Unfortunately, there are no effective treatments for most mitochondrial diseases [11,12], and so increased understanding of the cellular consequences of mtDNA damage is clearly imperative. *Saccharomyces cerevisiae* provides advantages as an experimental system in which to study mitochondrial dysfunction. For example, *S. cerevisiae* can survive the loss of mtDNA by generating sufficient ATP for viability by fermentation, and is therefore called a “petite-positive” yeast, based on historical nomenclature [13]. Upon additional perturbation of specific cellular functions and pathways, *S. cerevisiae* can become “petite-negative” and proliferate poorly or not at all following mtDNA loss. The petite-negative phenotype permits unbiased genetic screens and selections designed to reveal genes promoting or preventing fitness following mtDNA loss [14,15]. Consequently, findings apparently applicable across phylogeny to cells depleted of mtDNA, such as benefits provided by endomembrane system perturbation [16,17] and the need for a robust electrochemical potential (ΔΨ^mito^) across the mitochondrial IM [18–20], were first uncovered using budding yeast [14].

Since many biosynthetic and catabolic processes are localized to mitochondria, it is not surprising that mitochondrial abundance and function are responsive to the nutritional status of the cell [21–23]. Therefore, one avenue toward treatment of mitochondrial disorders may be the modulation of conserved, nutrient-sensing signaling pathways. Excitingly, recent findings obtained using yeast [24], worms [25], flies [26], and mammals [25,27] indicate that drugs and mutations affecting the Target of Rapamycin (TOR) pathway can alleviate the outcome of mitochondrial dysfunction, supporting the idea that a focus on signaling pathways controlled by nutrient levels is a rational approach toward treatment of mitochondrial disorders.

In this work, we have focused on the effects of glucose signaling on the outcome of mtDNA damage. We found that glucose restriction or inhibition of the glucose-sensing protein kinase A (PKA) pathway can lead to increased proliferation following mtDNA removal from *S. cerevisiae.* Interestingly, the benefits provided to cells lacking mtDNA by PKA inhibition require robust protein import from the cytosol. Increases in fitness coincide with a diminished transcriptional response to mtDNA damage, and we have discovered two new gene classes that are altered in their expression following mtDNA loss. Our work reveals that sugar sensation can control the physiology of cells lacking a mitochondrial genome.

## Results

### Reduction of PKA signaling can improve the fitness of cells lacking mitochondrial DNA

The PKA pathway controls the response to glucose in *S. cerevisiae* [28], and Pde2p is a phosphodiesterase that plays a dominant role in removing cyclic AMP (cAMP) and repressing PKA activity [29]. PKA hyperactivation by deletion of Pde2p or Ira2p leads to a loss of proliferation after mtDNA loss [15]. We speculated that PKA inhibition might, conversely, benefit cells lacking mtDNA. Toward this goal, we overexpressed Pde2p by transforming wild-type yeast with a high-copy plasmid containing the 2*μ* origin of replication and the *PDE2* gene. Indeed, Pde2p overproduction increased the proliferation of cells lacking functional mtDNA after ethidium bromide (EtBr) treatment [30] (or “*ρ*^-^ cells”) over that of *ρ*^-^ cells carrying an empty vector (Fig. 1A). Loss of functional mtDNA following EtBr treatment was confirmed by replica-plating to non-fermentable medium (S1 Fig.). Interestingly, the effects of PKA inhibition are dependent on yeast genetic background: while overexpression of Pde2p benefited *ρ*^-^ cells of the BY genetic background [31], *ρ*^-^ cells of the W303 background [32] did not increase their proliferation rate following Pde2p overexpression (S2 Fig.). Consequently, we continued to investigate the effects of PKA inhibition on mitochondrial dysfunction by taking advantage of the BY genetic background.

**Fig. 1:**
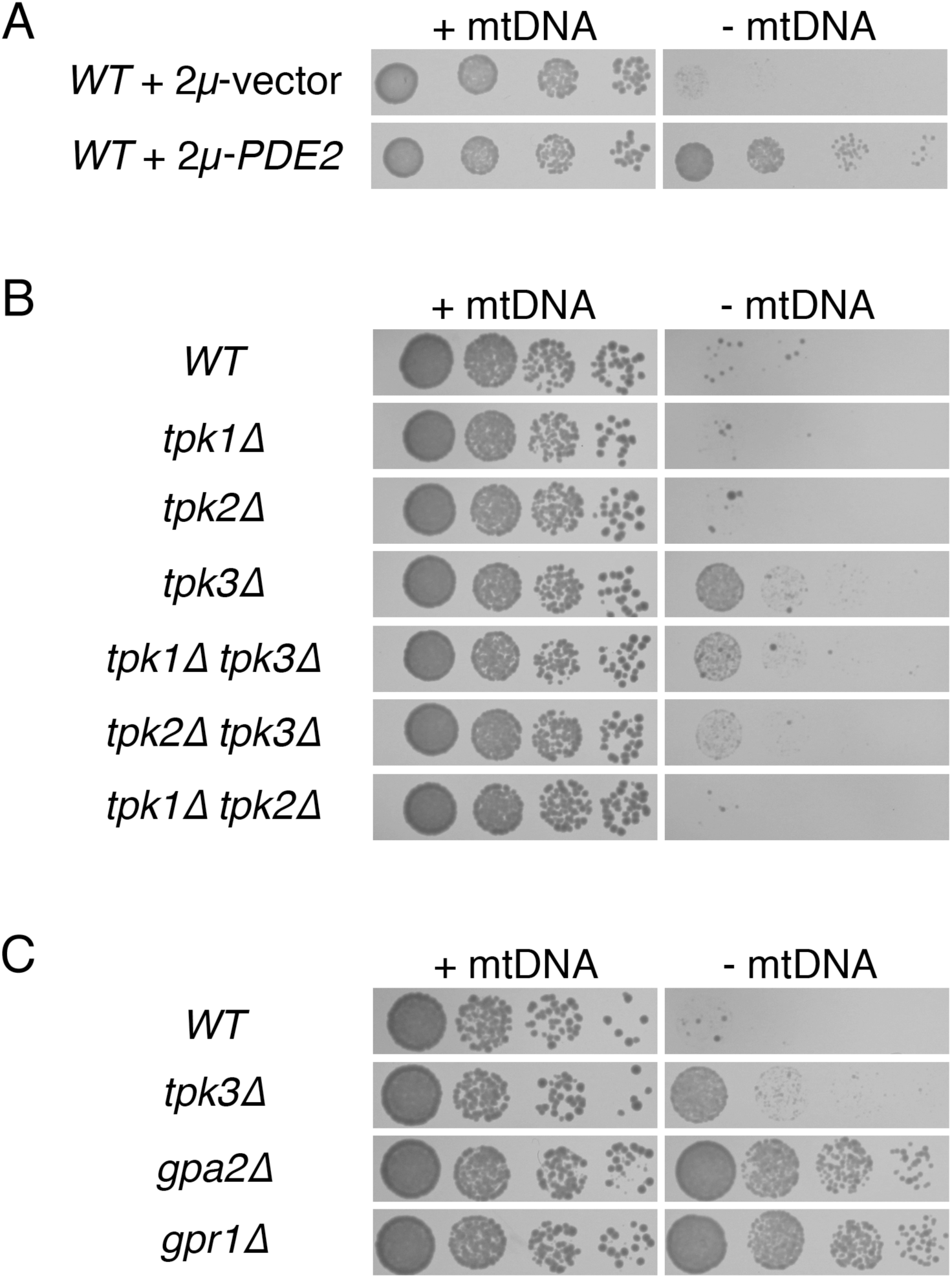
Decreased PKA activity can increase proliferation of cells lacking mtDNA. (A) Overexpression of cAMP phosphodiesterase Pde2p increases the fitness of cells lacking mtDNA. Strain BY4743 (*WT*) was transformed with empty, high-copy vector pRS426 or plasmid b89 (*2μ-PDE2*). Strains were tested for their response to mtDNA loss by incubation in selective medium lacking or containing 25 *μ*g/ml EtBr, with subsequent incubation on solid SC-Ura medium for 2d. (B) Lack of Tpk3p increases the fitness of cells lacking mtDNA. Strains BY4742 (*WT*), CDD884 (*tpk1Δ*), CDD885 (*tpk2Δ*), CDD886 (*tpk3Δ*), CDD908 (*tpk1Δ tpk3Δ*), CDD922 (*tpk2Δ tpk3Δ*), and CDD923 (*tpk1Δ tpk2Δ*) were tested for their response to mtDNA deletion with incubation on solid YEPD medium for 2d. (C) Cells deleted of Gpa2p or Gpr1p exhibit increased fitness after mtDNA deletion. Strains BY4742 (*WT*), CDD886 (*tpk3Δ*), CDD849 (*gpa2Δ*), and CDD850 (*gpr1Δ*) were treated as in (B).

Since PKA inhibition by Pde2p overexpression can increase the fitness of *ρ*^-^ cells, we asked whether any individual isoform of PKA might play a specific role in determining the outcome of mtDNA loss. Tpk1p, Tpk2p, and Tpk3p are the three PKA isoforms encoded by *S. cerevisiae* [28]. All three proteins act redundantly to allow cell proliferation, since *tpk1Δ tpk2Δ tpk3Δ* cells are normally inviable [33]. However, it is likely that each PKA isoform controls divergent cellular processes by phosphorylating its own specific set of target proteins [28,34]. We found that single deletion of Tpk3p increased the proliferation of *ρ*^-^ cells (Fig. 1B). Loss of the other PKA catalytic subunits, either singly or in combination, did not benefit cells lacking mtDNA. Interestingly, deletion of either Tpk1p or, more prominently, Tpk2p decreased the proliferation rate of *tpk3Δ ρ*^-^ cells, potentially suggesting a complex relationship between the PKA isoforms in which one PKA isoform might act upstream of another isoform in order to enact a signaling outcome.

PKA activity is promoted by Gpr1p, a G-protein coupled receptor, and Gpa2p, an associated G-protein α subunit [28,35]. We asked whether deletion of Gpr1p or Gpa2p would improve the fitness of *ρ*^-^ cells. Both *gpr1Δ ρ^-^* and *gpa2Δ ρ*^-^ cells proliferated more rapidly than isogenic *WT ρ*^-^ cells (Fig. 1C), further supporting a role for PKA activity in decreasing the fitness of cells lacking mtDNA. *ρ*^-^ cells lacking Tpk3p proliferate more slowly than *gpr1Δ ρ*^-^ cells and *gpa2Δ ρ*^-^ cells. This finding may suggest complex control of PKA isoforms by Gpr1p and Gpa2p beyond what is accessible by experiments using single and double Tpk deletion mutants. Alternatively this result may support an additional function for Gpr1p/Gpa2p outside of its role within the PKA pathway that is detrimental to proliferation of *ρ*^-^ cells.

### Glucose restriction benefits cells lacking mitochondrial DNA

Since a reduction in glucose sensation benefited cells lacking mtDNA, we next asked whether lowering glucose levels in the culture medium would similarly provide benefits to *ρ*^-^ cells. Indeed, lowering glucose from the standard concentration of 2% to either 0.5% or 0.2% resulted in improved proliferation of cells forced to lose mtDNA (Fig. 2A). Supporting the idea that glucose acts through the PKA pathway to determine the division rate of *ρ*^-^ cells, the benefits of the *gpr1Δ* or *gpa2Δ* mutations vanished upon medium with reduced glucose concentration (Fig. 2B).

**Fig. 2:**
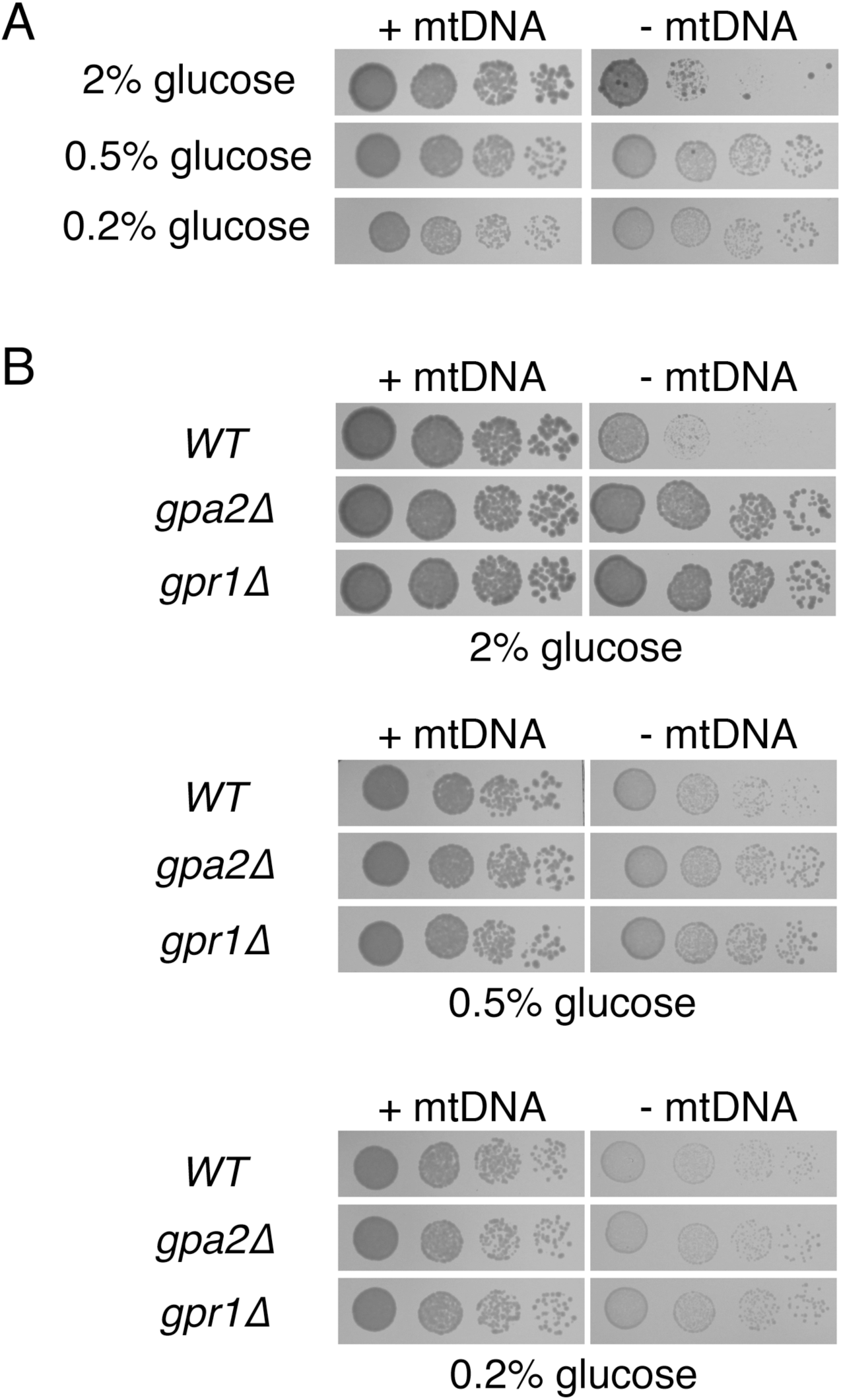
Glucose inhibits proliferation of cells deleted of mtDNA. (A) Decreasing glucose concentration leads to increased proliferation of cells lacking mtDNA. Strain BY4742 (*WT*) was cultured in YEPD medium containing 2%, 0.5%, or 0.2% glucose and tested for the response to mtDNA deletion. Cells were incubated for 3 d. (B) Proliferation of *ρ*^-^ cells by Gpa2p or Gpr1 p deletion is not improved further upon lowering the glucose concentration. Strains BY4742 (*WT*), CDD849 (*gpa2Δ*), and CDD850 (*gpr1Δ*) were treated as in (A), yet incubated on solid medium for 2 d.

### The transcriptional response to mtDNA loss is diminished upon PKA inhibition

A transcriptional response in the nucleus is activated following mtDNA loss. Most prominently, genes activated by an iron deprivation response (IDR) can be highly induced by mtDNA deletion [24,36]. Furthermore, the pleiotropic drug resistance (PDR) pathway can be induced by mtDNA loss [37], as is the expression of several tricarboxylic acid cycle enzymes, through activation of the yeast retrograde (RTG) pathway [38,39]. Interestingly, this transcriptional response is often abated under conditions where *ρ*^-^ cells exhibit greater fitness [36,40]. By next-generation sequencing of cellular transcripts (RNA-seq), we investigated whether the transcriptional response to mtDNA deletion is altered in cells with reduced PKA activity. Toward this goal, we compared the gene expression of *ρ^+^* cells to that of *ρ*^-^ cells either overexpressing or not overexpressing Pde2p. We also overexpressed Tip41p in *ρ*^-^ cells as a positive control, since perturbation of the arm of the TORC1 pathway in which Tip41p functions ameliorates the gene expression changes precipitated by mtDNA loss [24].

We first assessed the expression level of *PDE2* and *TIP41* transcripts in our samples, and we also investigated potential selective pressure against overexpression of these genes. We noted that *ρ*^-^ cells overexpressing Pde2p do so with *PDE2* transcript levels averaging nearly four-fold higher than *ρ*^-^ cells incorporating an empty vector, while *ρ*^-^ cells overexpressing Tip41p contain *TIP41* transcript levels 12-fold higher than *ρ*^-^ cells containing an empty vector (S1 Table). High-copy plasmids used for gene overexpression contained the *URA3* gene as a selectable marker. When compared to *URA3* expression within *ρ*^-^cells containing an empty 2*μ*-*URA3* vector, *ρ*^-^cells overexpressing Tip41p contain 61% less *URA3* transcript, and *ρ*^-^ cells overexpressing Pde2p possess 86% less *URA3,* suggesting that cells that express Tip41p or Pde2p at extreme levels might be lost from the population. These findings are congruent with the essential nature of PKA signaling [33] and of signaling through the Tap42p-dependent arm of the TORC1 pathway [41].

As expected, genes within the IDR pathway (Fig. 3A), the PDR pathway (Fig. 3B), and the RTG pathway (Fig. 3C) were significantly induced by mtDNA deletion. Consistent with improved fitness of cells lacking mtDNA following PKA inhibition, transcription of genes within all three of these pathways was diminished in *ρ*^-^ cells overexpressing Pde2p (Figs. 3A-3C). Excitingly, our analysis revealed that *ARG1, ARG3, ARG4, ARG5,6, ARG7, CPA1,* and *CPA2,* all encoding genes involved in arginine biosynthesis [42] were induced upon mtDNA loss (q <0.05, S1 Table). We focused our attention upon *ARG1, CPA2,* and *ARG3,* which were induced at least three-fold by mtDNA loss, and we found that the activation of these transcripts in *ρ*^-^ cells was reduced upon PKA inhibition (Fig. 3D). Finally, we encountered a significant down-regulation of *ARO9* and *ARO10* following mtDNA deletion (Fig. 3E). These two genes play roles in aromatic amino acid catabolism [43]. The downregulation of these two targets upon mtDNA deletion was also alleviated by PKA inhibition. Together, our data indicate that changes to a nuclear transcription program that is associated with mtDNA damage can be lessened by reduced PKA activity.

**Fig. 3:**
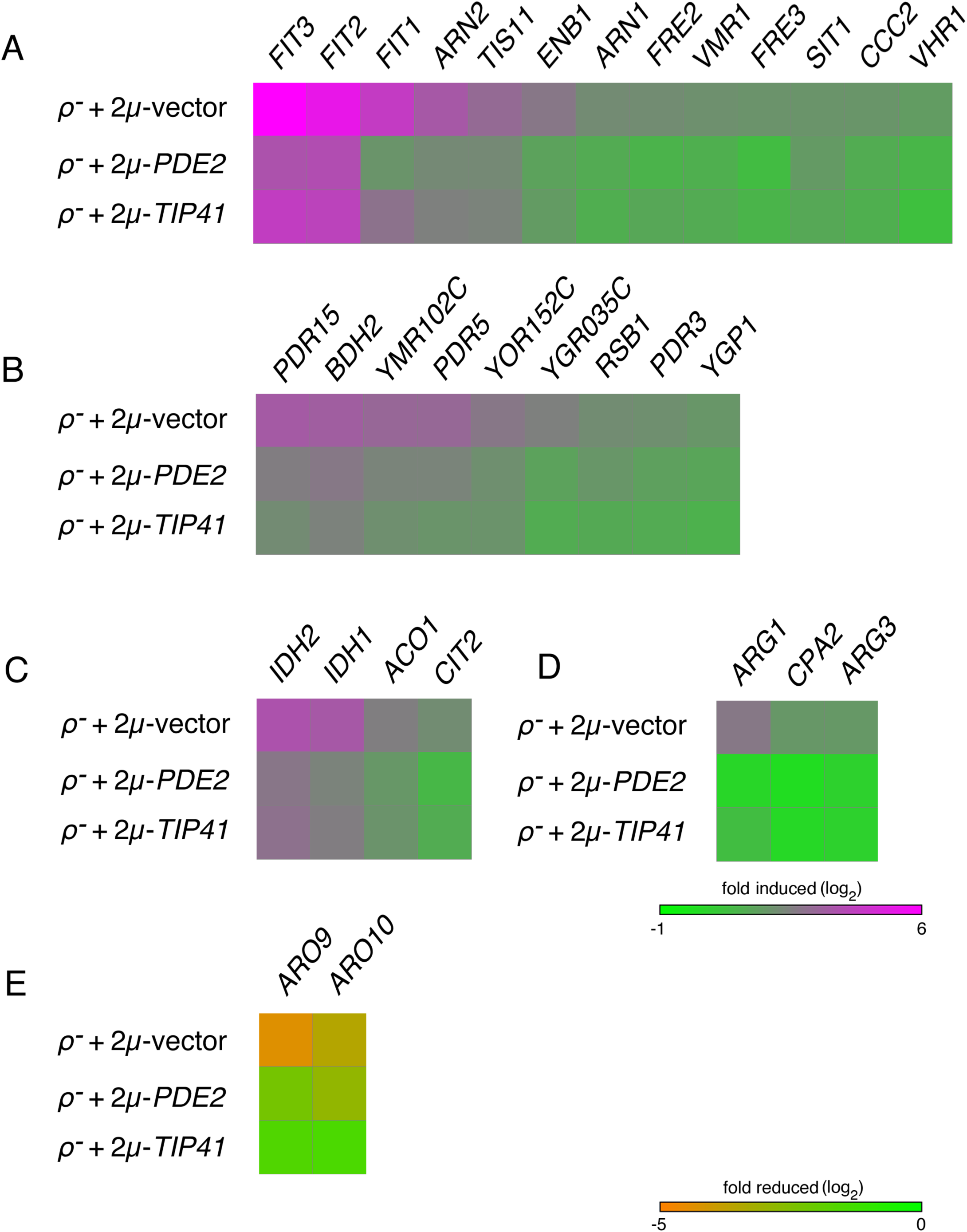
Overexpression of Pde2p can diminish the transcriptional response to mtDNA deletion. (A) IDR target genes activated by mtDNA loss are attenuated upon Pde2p overexpression. Wild-type strain BY4743 transformed with vector pRS426, plasmid b89 (pRS426-PDE2), or plasmid M489 (pRS426-*TIP41*) was treated with EtBr for 24 hr to force mtDNA loss. Gene expression levels were determined by next-generation sequencing and normalized to BY4743 *ρ^+^* cells harboring vector pRS426. Genes selected for analysis were activated more than three-fold in *ρ*^-^ cells expressing vector pRS426 over *ρ^+^* cells expressing the same plasmid, were statistically significant upon comparison of these two conditions (q < 0.05), and were listed as IDR targets in [124] or [125]. (B) Genes activated by the PDR pathway in *ρ*^-^ cells are reduced in expression by Pde2p overexpression. Analysis was performed as in (A), except genes selected for analysis were identified as PDR pathway targets in [126] or [127]. (C) Genes activated by the RTG signaling pathway in cells lacking a mitochondrial genome are decreased in expression by Pde2p overproduction. Analysis was performed as in (A), with RTG pathway targets provided by [40]. (D) Arginine biosynthesis genes are upregulated upon mtDNA loss, but this response is reduced upon PKA inhibition. Analysis was performed as in (A), with arginine biosynthesis genes reported by [121]. (E) Two genes involved in aromatic amino acid breakdown and reduced in expression following mtDNA loss are recovered in expression when Pde2p is overexpressed. Analysis was performed as in (A). *ARO9* and *ARO10* were selected after a more than three-fold (q < 0.05) reduction in expression when comparing *ρ*^-^ cells to *ρ^+^* cells containing vector pRS426. Quantitative expression data can be found in S1 Table. Data are visualized using [128].

### Import of a mitochondria-directed protein is not augmented by PKA inhibition in cells lacking mtDNA

Upon deletion of mtDNA, ΔΨ^mito^ is reduced [44]. Consequently, protein import of essential, nucleus-encoded proteins through the voltage-gated IM translocons is curtailed [36,45]. Deletion of protein phosphatases acting within the TORC1 pathway [24] or blockade of vacuolar acidification [16] were previously demonstrated to reverse the effects of mtDNA damage and to increase the mitochondrial localization of an *in vivo* reporter of mitochondrial protein import. Therefore, we asked whether reduction of PKA activity might similarly result in heightened mitochondrial protein import in cells lacking mtDNA. Toward this goal, we used an *in vivo* reporter of mitochondrial protein import that consists of the first 21 amino acids of Cox4p fused to a fluorescent protein [36]. As previously reported, this protein localized to mitochondria in *ρ^+^* cells, but was poorly imported into the mitochondria of *ρ*^-^ cells (Fig. 4A). Cox4p(1-21)-GFP was expressed in *WT, gpa2Δ* and *gpr1Δ* cells, yet in spite of the clear benefit to cellular fitness provided by deletion of these drivers of PKA signaling upon mtDNA loss, no redistribution of Cox4p(1-21)-GFP to *ρ^-^* mitochondria of *gpa2Δ* or *gpr1Δ* cells was detected (Fig. 4B and 4C). As a positive control, we included cells lacking subunit Vma2p of the vacuolar proton pumping ATPase. Cells deleted of Vma2p have been demonstrated to exhibit improved protein import following mtDNA loss [16], and mitochondrial localization into *ρ^-^* mitochondria in this nuclear background was confirmed. Cox4p(1-21)-GFP was also not re-localized to *ρ^-^* mitochondria in cells overexpressing Pde2p, but an empty 2*μ*-vector containing the *URA3* marker led to increased Cox4p(1-21)-GFP localization to mitochondria lacking mtDNA, making interpretation of any microscopy results using the same vector containing *PDE2* difficult to interpret (unpublished results). Our findings do not support augmented protein import into *ρ^-^* mitochondria upon reduction of PKA activity, potentially distinguishing those mutants with diminished PKA activity from other mutants previously demonstrated to improve the fitness of cells deleted of the mitochondrial genome.

**Fig. 4:**
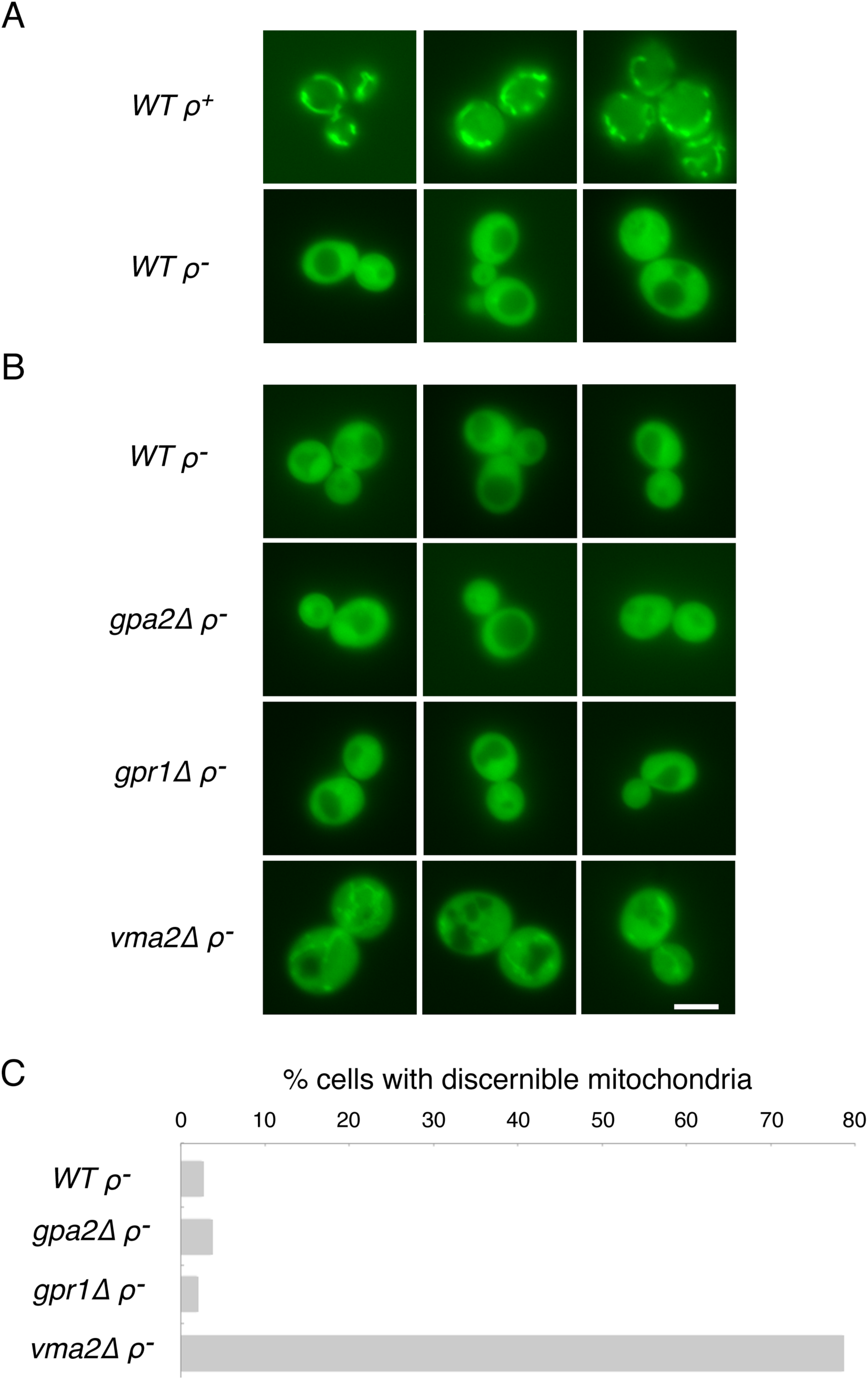
Cells lacking the glucose-sensing proteins Gpa2p or Gpr1p do not increase the localization of a mitochondria-targeted fluorescent protein to mitochondria. (A) Strain BY4742 (*WT*) carrying pHS12 and either harboring or lacking mtDNA was examined by fluorescence microscopy to demonstrate the mitochondrial location of Cox4p(1–21)-GFP in *ρ^+^* cells and its cytosolic location *ρ*^-^ cells. Mitochondria are visible in >99% of the cells within a *ρ^+^* culture [16] (B) Strains BY4742 (*WT*), CDD849 (*gpa2Δ*), CDD850 (*gpr1Δ*), and CDD496 (*vma2Δ*) carrying pHS12 and deleted of mtDNA were examined by fluorescence microscopy, with representative images displayed. Scale bar, 5*μ*m. (C) These cultures were scored, blind to genotype, for localization of Cox4p(1–21)-GFP to mitochondria (n > 200 cells).

### Several well-characterized proteins and pathways controlled by PKA signaling are not individually responsible for the beneficial effects of PKA inhibition on cells lacking mtDNA

Lack of PKA signaling activates pathway responsive to a stressful cellular environment. Prominent transcription factors activated following PKA inhibition and driving stress resistance include Gis1p [46] and the paralogous Msn2 and Msn4 proteins [47,48]. We tested whether transcriptional activation of these factors may be solely responsible for the beneficial effects provided by PKA inhibition, but both *gis1Δ ρ*^-^ cells (Fig. 5A) and *msn2Δ msn4Δ ρ*^-^cells (Fig. 5B) responded positively to Pde2p overexpression. The Rim15 kinase integrates the activities of several signaling pathways, including PKA, to control Gis1p and Msn2p/Msn4p [49], but like cells deleted of its targets, *rim15Δ* mutants lacking mtDNA are benefited by PKA inhibition (Fig. 5C).

**Fig. 5:**
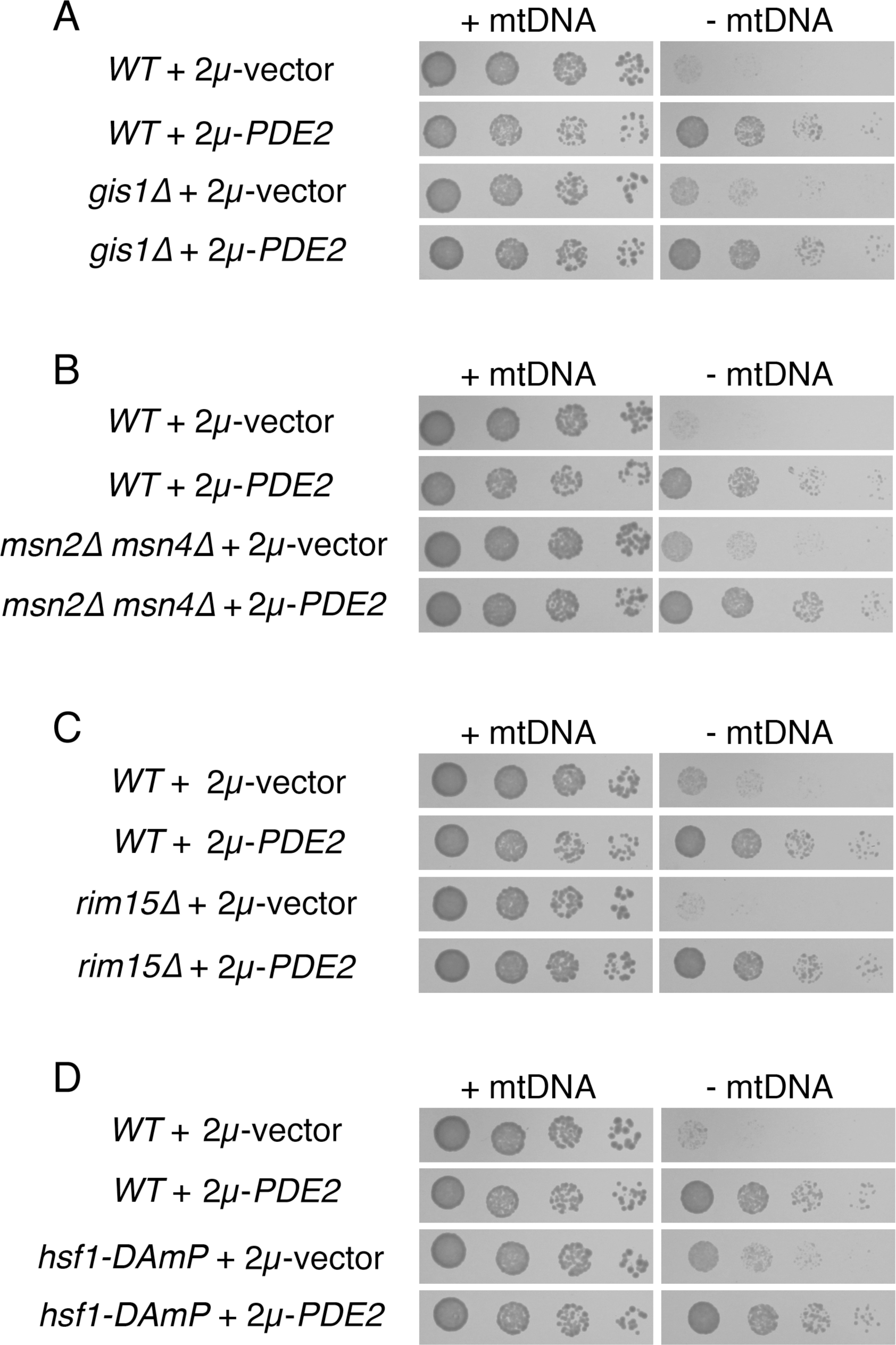
Several transcription factors driving stress resistance following PKA inhibition are not individually responsible for the benefits provided by Pde2p overexpression to cells lacking mtDNA. (A) Cells lacking Gis1p and mtDNA are increased in proliferation upon Pde2p overexpression. Strains BY4742 (*WT*) and CDD801 (*gis1Δ*) were treated as in Fig. 1A. (B) Cells lacking both Msn2p and Msn4p exhibit increased fitness following mtDNA loss upon Pde2p overexpression. Strains BY4741 (*WT*) and CDD838 (*msn2Δ msn4Δ*) were treated as in Fig. 1A. (C) The Rim15 kinase is not required in order for Pde2p overexpression to benefit *ρ*^-^ cells. Strains CDD463 (*WT*) and CDD841 (*rim15Δ*) were treated as in Fig. 1A. (D) A potential reduction of Hsf1p function does not prevent increased *ρ^-^* cell fitness upon overexpression of Pde2p. Strains BY4741 (*WT*) and CDD910 (*hsf1-DAmP*) were treated as in Fig. 1A.

Since the essential transcription factor Hsf1p, which controls a host of stress-induced genes [50], can be inhibited via PKA signaling [51], we wondered whether induction of Hsf1p targets may be relevant to the fitness of *ρ*^-^ cells overexpressing Pde2p. However, a strain with potentially attenuated expression of Hsf1p due to disruption of the *HSF1* 3' untranslated region [52] was responsive to *2μ-PDE2* after mtDNA loss (Fig. 5D), and in fact *hsf1-DAmP ρ*^-^ cells divided more rapidly than expected when compared to *HSF1 ρ*^-^ cells (Fig. 5D and unpublished results). Moreover, our RNA-seq analysis suggested no appreciable activation of several targets containing Hsf1p binding sites [53] upon PKA signal reduction in *ρ*^-^cells, including *SSA1, SSA2, SSA3, SSA4, HSP78, BTN2, HSP104,* and *SIS1* (S1 Table). Therefore, it is unlikely that Hsf1p activation could be solely responsible for the benefits provided by PKA reduction to *ρ*^-^ cells.

Two chaperones, Hsp12p and Hsp26p, are prominently induced by a reduction in PKA signaling through Msn2p/Msn4p, Gis1p, and Hsf1p [46,51,54-56]. Indeed, *HSP12* and *HSP26* transcripts are increased greater than 7-fold and 4-fold, respectively, in *ρ*^-^ cells harboring a *2μ-PDE2* plasmid when compared to *ρ*^-^ cells carrying an empty vector (Fig. 6A). We asked whether an increase in the production of Hsp12p or Hsp26p might be the mechanism by which reduced PKA activity benefits cells lacking mtDNA. Both *hsp12Δ ρ*^-^ cells and *hsp26Δ ρ*^-^ cells increased their proliferation rate in response to Pde2p overexpression (Fig. 6B), indicating that neither Hsp12p nor Hsp26p were the lone mediators of the response to PKA reduction. These findings are consistent with the results obtained using strains lacking various transcriptional regulators of Hsp12p and Hsp26p expression.

**Fig. 6:**
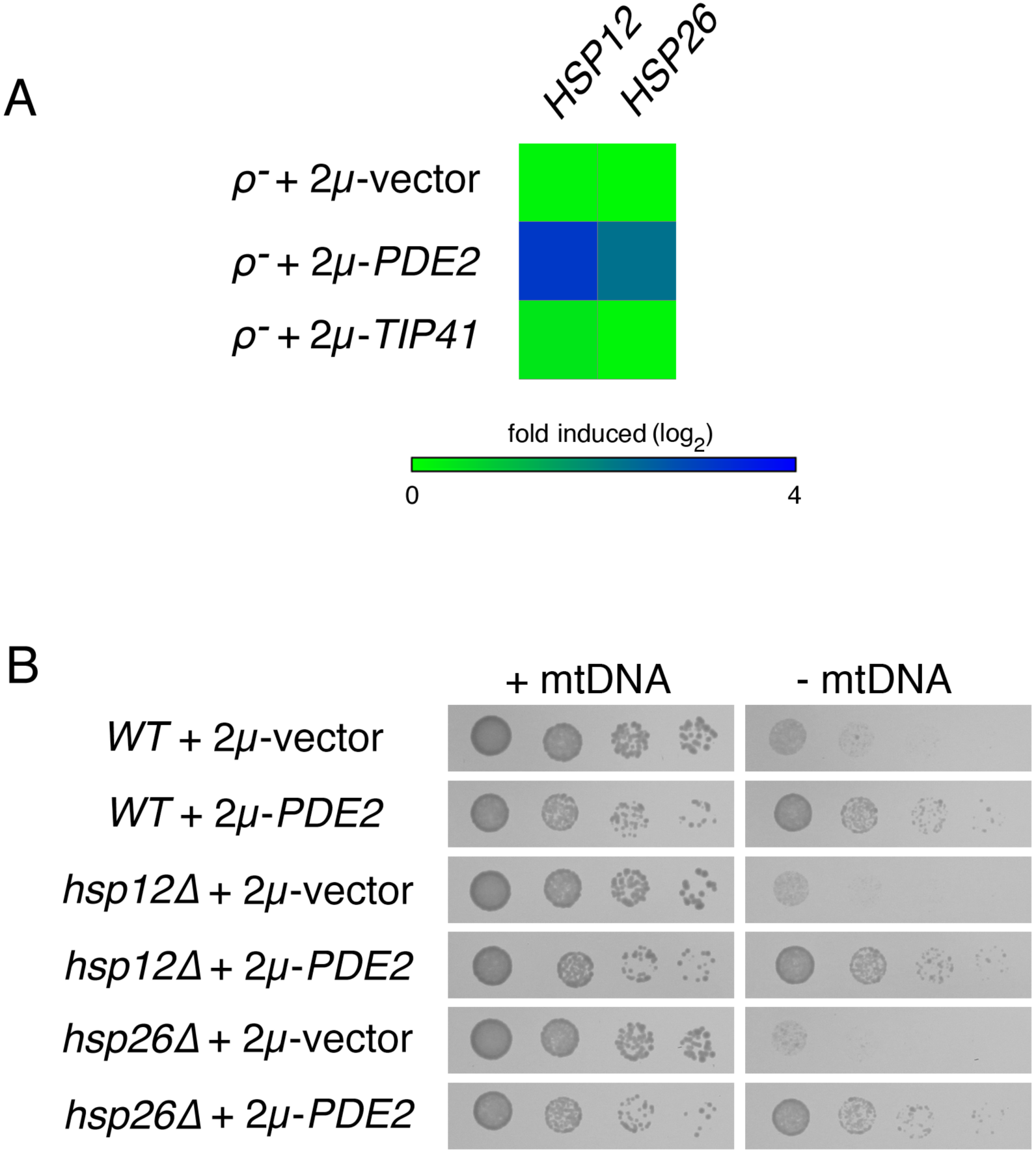
Chaperones Hsp12p and Hsp26p are not required for increased proliferation of cells lacking mtDNA upon Pde2p overexpression. (A) *HSP12* and *HSP26* transcripts are overexpressed upon PKA inhibition in *ρ*^-^ cells. *HSP12* and *HSP26* transcript abundance within *ρ*^-^ cells upon overexpression of Pde2p, overproduction of Tip41 p, or upon maintenance of empty vector pRS426 were normalized to gene expression in *ρ^+^* cells carrying an empty vector. (B) Neither Hsp12p nor Hsp26p are individually responsible for the benefits provided by Pde2p overexpression to cells deleted of mtDNA. Strains CDD463 (*WT*), CDD542 (*hsp12Δ*), CDD534 (*hsp26Δ*) were treated as in Fig. 1A.

PKA inhibition not only activates genes involved in stress resistance, but also leads to inhibition of cytosolic protein synthesis and cell proliferation. In parallel with the TORC1-regulated Sch9 protein, PKA phosphorylates and inhibits the highly related transcriptional repressors Dot6p and Tod6p to promote the transcription of proteins involved in cytosolic translation [57–59]. Moreover, reduction of cytosolic translation by mutation or pharmacological inhibition increases the fitness of yeast lacking mtDNA [15,60–62]. We asked whether *ρ^-^* mutants lacking both Dot6p and Tod6p would still respond to PKA inhibition, and we found that *dot6Δ tod6Δ ρ^-^* proliferation was enhanced by an increased dosage of Pde2p (Fig. 7A). Therefore, PKA inhibition does not benefit cells lacking mtDNA solely by promoting Dot6p and Tod6p repression of transcriptional targets.

**Fig. 7:**
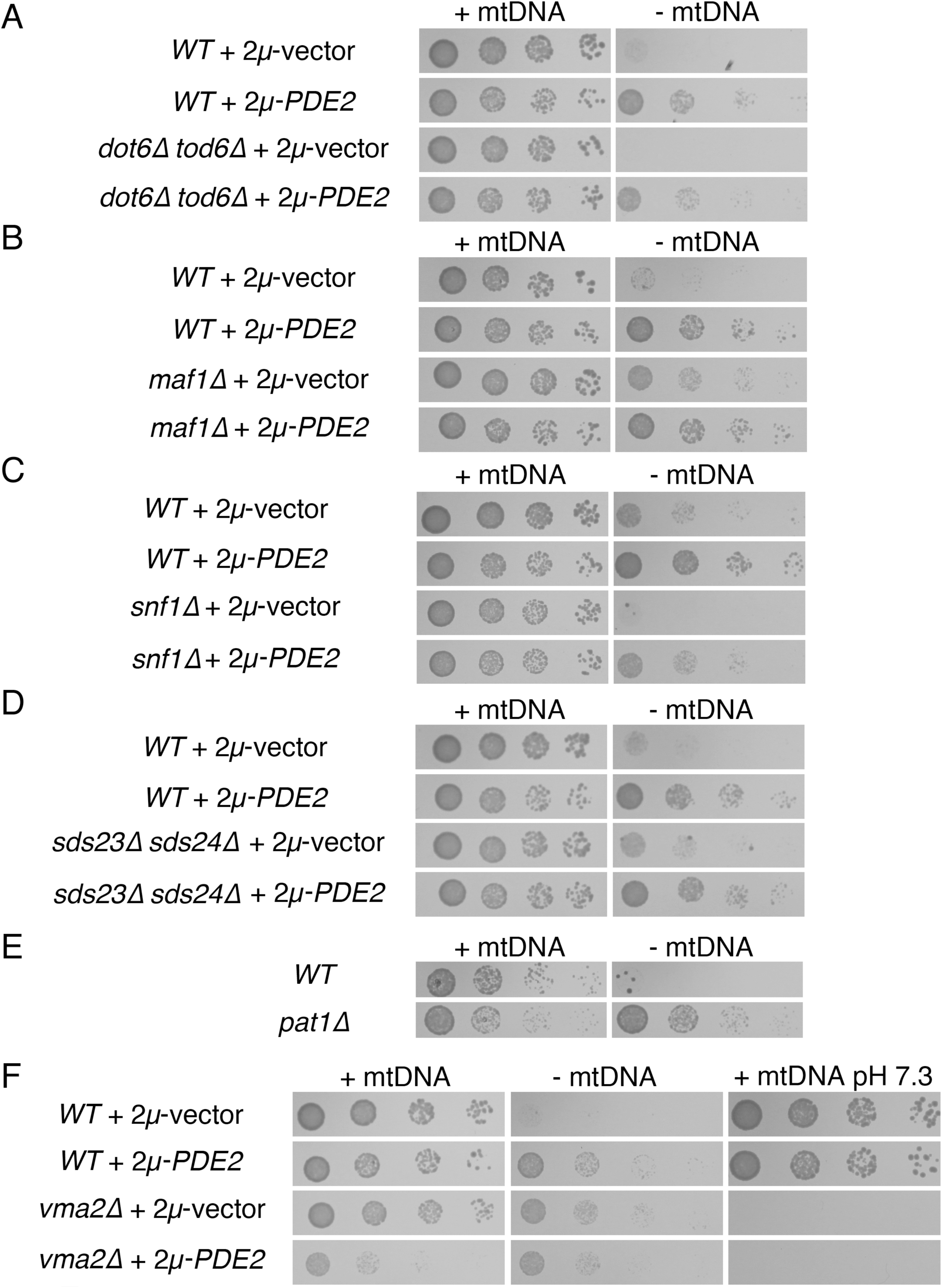
Several cellular processes and signaling pathways controlled by PKA activity are not individually responsible for the outcome of PKA inhibition for cells deleted of mtDNA. (A) Repression of transcriptional targets of Dot6p and Tod6p is not the sole mechanism by which high-copy Pde2p benefits *ρ*^-^ cells. Strains CDD289 (*WT*) and CDD567 (*dot6Δ tod6Δ*) were treated as in Fig. 1A. (B) Repression of Maf1 p targets is not required in order for Pde2p overexpression to benefit cells lacking mtDNA. Strains BY4741 (*WT*) and CDD928 (*maf1Δ*) were treated as in Fig. 1A. (C) Activity of the Snf1 kinase is not required in order for Pde2p overproduction to increase proliferation of cells deprived of a mitochondrial genome. Strains CDD463 (*WT*) and CDD604 (*snf1Δ*) were treated as in Fig. 1A. (D) Deletion of Sds23p and Sds24p does not prevent overexpression of Pde2p from increasing the division rate of cells lacking mtDNA. Strains BY4741 (*WT*) and CDD921 (*sds23Δ sds24Δ*) were treated as in Fig. 1A. (E) Cells lacking P-body component Pat1p exhibit increased fitness after mtDNA loss. Strains BY4742 (*WT*) and CDD879 (*pat1Δ*) were treated as in Fig. 1B, except *ρ^+^* cells were incubated on solid YEPD medium for 1d, while *ρ*^-^ cells were incubated for 2 d. (F) PKA inhibition by Pde2p overexpression is unlikely to significantly affect V_1_V_o_-ATPase assembly. Strains BY4742 (*WT*) and CDD496 (*vma2Δ*) were treated as in Fig. 1A. *ρ^+^* cultures were additionally plated to SC-Ura medium buffered to pH 7.3 using 100 mM HEPES-KOH, pH 7.5 and incubated for 3 d.

Maf1p also contributes to the control of cytosolic translation via its regulation of RNA polymerase III [63,64]. PKA has been reported to phosphorylate and negatively regulate Maf1p [65]. We tested whether deletion of Maf1p would prevent PKA inhibition from increasing the proliferation of *ρ*^-^ cells, and we found that *maf1Δ ρ*^-^ cells proliferated more rapidly upon overproduction of Pde2p (Fig. 7B). These results indicate that PKA inhibition cannot reverse the effects of mitochondrial dysfunction solely through activation of Maf1p.

Previously, we found that Snf1p, the catalytic subunit of the AMP-sensitive protein kinase ortholog of *S. cerevisiae,* can play a definitive role in the outcome of mtDNA loss [24]. Moreover, the Sip1 protein, thought to direct Snf1p toward specific substrates, is altered in its subcellular location following PKA inhibition [66]. Therefore, we tested whether cells lacking Snf1p or Sip1p could exhibit increased proliferation following mtDNA loss if Pde2p was overexpressed. However, the petite-negative phenotype of *snf1Δ ρ*^-^ cells was rescued by PKA inhibition (Fig. 7C), and *sip1Δ ρ*^-^ cells proliferated more quickly upon Pde2p overproduction (unpublished results), indicating that Snf1p activation is not the sole mechanism by which PKA inhibition benefits cells deprived of mtDNA.

The paralogous Sds23 and Sds24 proteins may act in opposition to PKA signaling [67] and are also likely to inhibit type 2A and 2A-like phosphatases [68,69]. It was previously shown that *sds24Δ* mutants are sensitive to mtDNA loss [24], and so we examined how cells lacking both Sds23p and Sds24p and mtDNA might respond to Pde2p overexpression. Surprisingly, a newly generated *sds24Δ* strain proliferated as rapidly as a *WT* strain following mtDNA loss on YEPD medium (S3 Fig.), and a *sds23Δ sds24Δ ρ^-^* strain can proliferate more rapidly than a *WT ρ^-^* strain on rich medium (unpublished results). Deletions and auxotrophic markers were confirmed in previous and current strains, suggesting that previous results might be a consequence of a background mutation inherited from a parental strain obtained from the deletion mutant collection [70]. Regardless, overexpression of Pde2p could provide increased fitness to *sds23Δ sds24Δ ρ*^-^cells (Fig. 7D), indicating that PKA inhibition does not benefit *ρ*^-^cells solely through control of Sds23p or Sds24p.

Processing Bodies, or “P-bodies,” can form when cells encounter stressful conditions, and P-bodies are thought to be storage sites for mRNA transcripts not undergoing translation and potentially destined for degradation [71]. Interestingly, Tpk3p can be localized to P-bodies, and P-bodies form more quickly following Tpk3p deletion [72]. Furthermore, P-body formation can be prevented by PKA phosphorylation of the conserved Pat1 protein [73], a factor that normally promotes P-body generation [74]. Therefore, we hypothesized that PKA inhibition could increase *ρ^-^* cell fitness by increasing P-body formation or functionality. However, we found that *pat1Δ ρ*^-^ cells exhibit a considerably higher division rate than *WT* cells (Fig. 7E), indicating that modulation of Pat1p cannot be wholly responsible for the benefits cells provided by PKA inhibition to cells lacking mtDNA.

Intriguing functional and physical connections exist between mitochondria and vacuoles [16,17,75–77]. Importantly, increasing vacuolar pH has been found to improve the fitness of yeast and protist cells lacking mtDNA. Some evidence suggests that PKA can modulate vacuolar V_1_V_o_-ATPase assembly [78]. Overexpression of Pde2p did not seem to provide a highly evident and reproducible proliferation benefit to *ρ*^-^ cells already lacking Vma2p, potentially suggesting a role for V_1_V_o_-ATPase disassembly in determining the outcome of PKA signaling in *ρ*^-^ cells (Fig. 7F). However, while cells with reduced or absent V_1_V_o_-ATPase activity, such as the *vma2Δ* mutant, manifest defective proliferation on alkaline medium [79,80], *WT ρ^+^* cells overexpressing Pde2p exhibit no proliferation defect at all on alkaline medium. This finding argues against PKA pathway inhibition modulating vacuolar pH to determine the proliferation rate of *ρ*^-^ cells.

### Overexpression of Pde2p can suppress the petite-negative phenotype of several mutants characterized by defective mitochondrial function

While *S. cerevisiae* can typically survive mtDNA loss, mutation of a number of nuclear genes can lead to severe proliferation defects, or a “petite-negative” phenotype [13], following mtDNA deletion. For example, mutants lacking the full function of the i-AAA protease, which is involved in protein quality control at the IM, are affected by mtDNA deletion [15,81,82]. We tested whether the petite-negative phenotype of cells lacking Mgr1p or Mgr3p, accessory subunits of the i-AAA protease, could be suppressed by Pde2p overexpression. Indeed, *mgr1Δ ρ^-^* and *mgr3Δ ρ*^-^ cells could proliferate following mtDNA loss if carrying the *2μ-PDE2* plasmid (Fig. 8A). Next, we examined the ability of PKA inhibition to rescue the petite-negative phenotype of cells deleted of Mgr2p, a component of the TIM23 complex that controls lateral segregation of proteins into the IM [83], and we found that *mgr2Δ ρ*^-^ cells overexpressing Pde2p were viable (Fig. 8B). Furthermore, we tested whether cells lacking Phb1p, required for formation of the prohibitin complex of the IM, could proliferate following mtDNA loss if PKA were inhibited. The prohibitin complex may scaffold the assembly of IM protein complexes [84]. Indeed, overexpression of Pde2p was able to suppress the petite-negative phenotype of *phb1Δ* mutants.

**Fig. 8:**
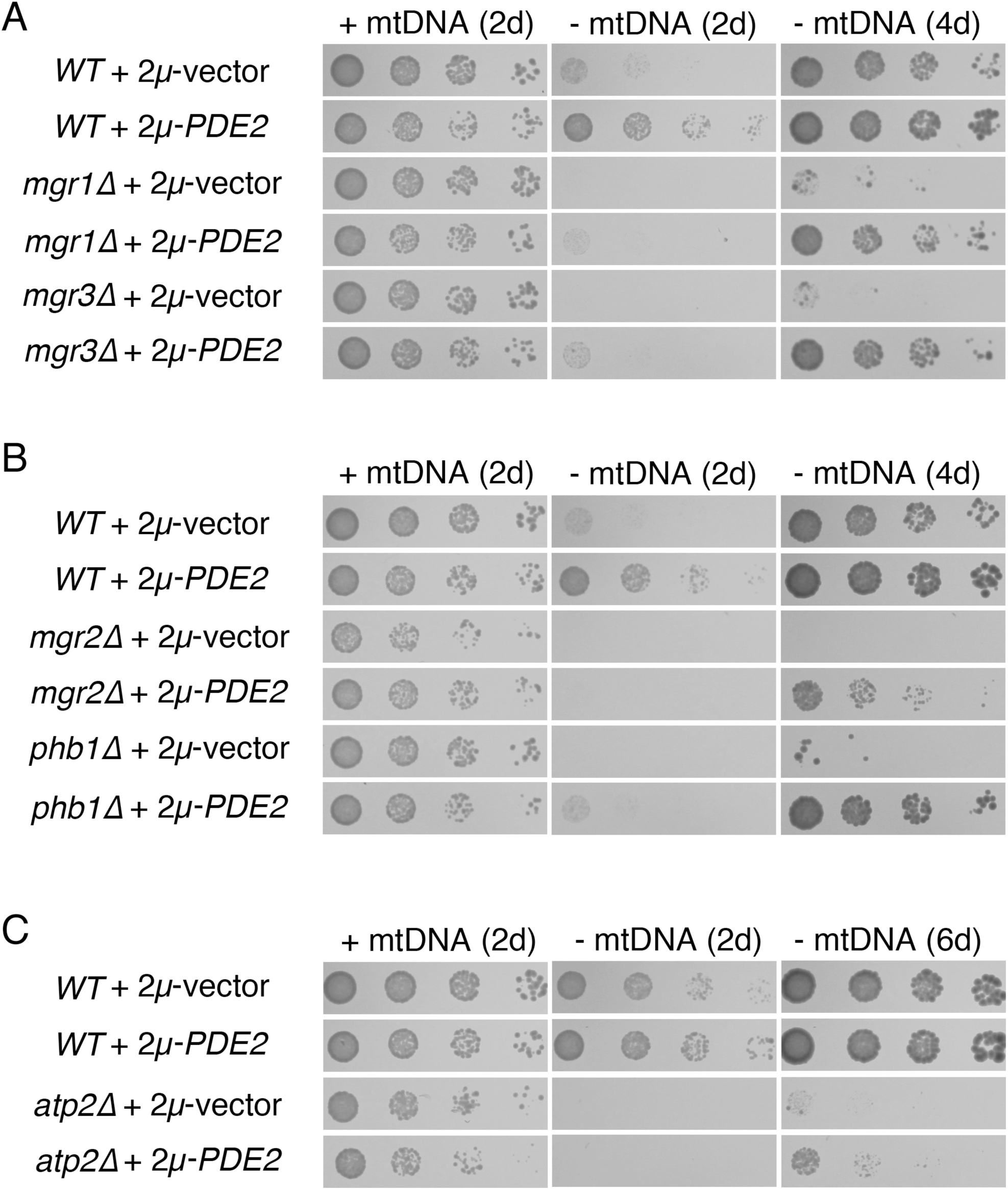
Overexpression of Pde2p can rescue the petite-negative phenotype of several mutants defective for mitochondrial function. (A) High-copy Pde2p can allow mutants deficient in activity of the i-AAA protease to remain viable following mtDNA loss. Strains BY4741 (*WT*), CDD13 (*mgr1Δ*), and CDD15 (*mgr3Δ*) were treated as in Fig. 1A, with additional incubation of *ρ*^-^ cells to 4 d in order to demonstrate suppression of the petite-negative phenotype. (B) PKA inhibition by Pde2p overexpression can rescue the petite-negative phenotype of mutants deficient in mitochondrial protein import and assembly. Strains BY4741 (*WT*), CDD11 (*mgr2*Δ), and CDD17 (*phb1Δ*) were treated as in (A). (C) Overexpression of Pde2p allows cells lacking mtDNA to proliferate in the absence of F_1_-ATPase activity. Strains CDD463 (*WT*) and CDD215 (*atp2Δ*) were treated as in Fig. 1A, with further incubation of *ρ*^-^ cells to 6d.

The F_1_ portion of the ATP synthase plays a major role in generating the essential ΔΨ^mito^ of *ρ^-^* mitochondria through a poorly understood electrogenic circuit thought to require the F_1_ sector’s ability to hydrolyze ATP [14,18,20,85–87]. The F_1_ sector performs this role even in the absence of a functional F_O_ sector of the ATP synthase [88], which is partially encoded by the mitochondrial genome. Interestingly, the *atp2Δ* mutant, which lacks F_1_ activity, was also rescued following mtDNA loss by attenuation of PKA (Fig. 8C).

The generation of ΔΨ^mito^ within *ρ^-^* mitochondria also requires Aac2p, the major mitochondrial ATP/ADP antiporter of the IM, in order to exchange ADP produced in the matrix by the F_1_ sector with more negatively charged ATP [44,86,89]. We generated an *aac2Δ* strain of mixed genetic background in order to acquire a functional *SAL1* allele, which is required for *aac2Δ* viability even when mtDNA is present [90]. In contrast to the petite-negative mutants assayed above, cells lacking both Aac2p and mtDNA were not rescued by Pde2p overexpression (unpublished results). Such a result might be anticipated: sufficient ATP import into mitochondria is expected to be required by *ρ*^-^ cells even outside of the context of ΔΨ^mito^ generation, since matrix ATP is necessary in order for matrix-localized chaperones to drive the essential mitochondrial protein import process [91,92]. The petite-negative phenotype of cells lacking Aac2p has, to our knowledge, never been suppressed.

### The mitochondrial protein import receptor Tom70 is required for cells lacking mtDNA to receive the benefits provided to by PKA inhibition

Tom70p, an outer membrane receptor playing a prominent role in the import of hydrophobic proteins with internal targeting information into mitochondria, can be required for viability following mtDNA loss [93]. Intriguingly, *tom70Δ ρ*^-^ cells were not rescued by overexpression of Pde2p (Fig. 9A), while, as previously demonstrated, high-copy Tip41p was able to rescue the petite-negative phenotype of *tom70Δ* mutants.

**Fig. 9:**
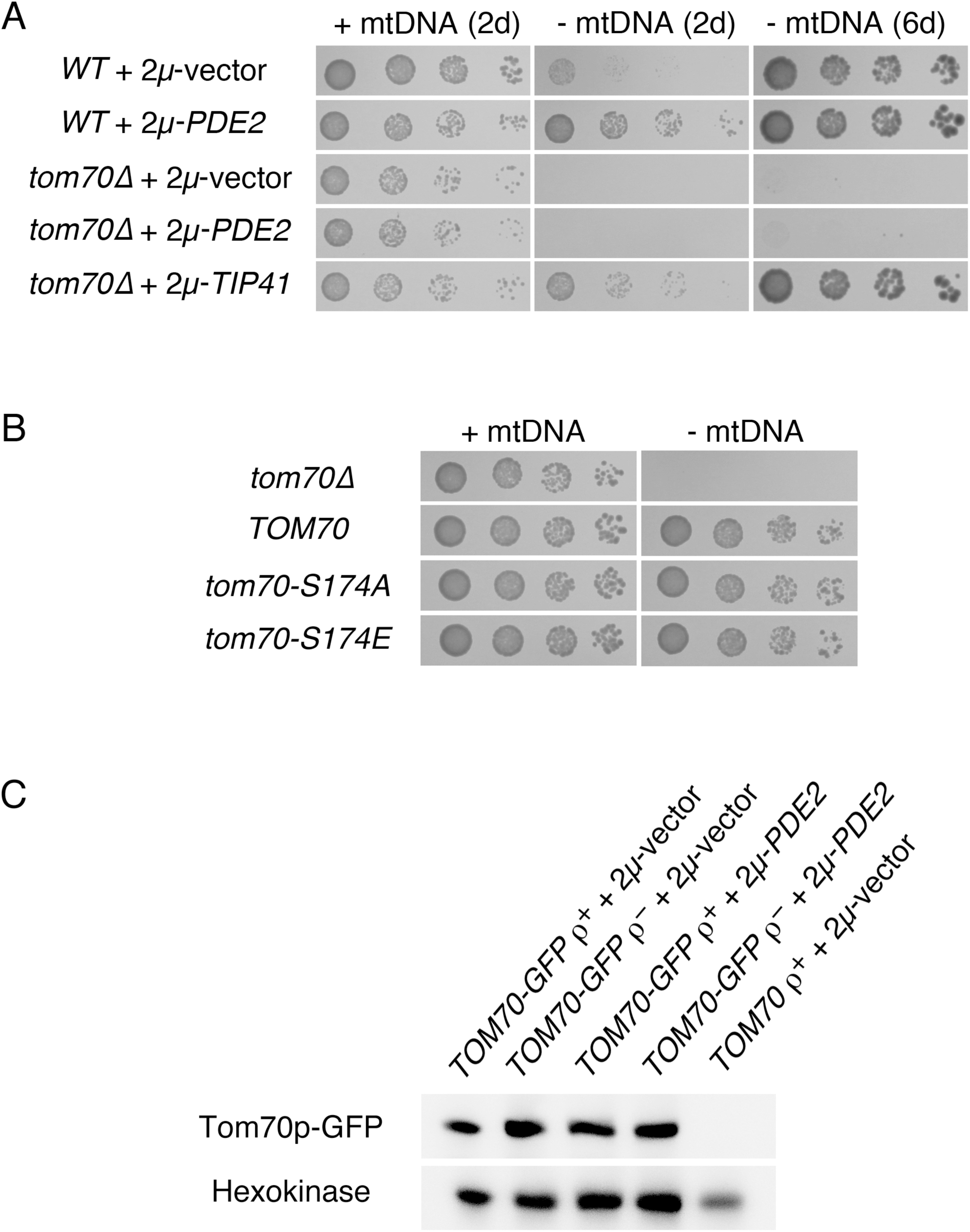
The outer membrane protein receptor Tom70 is required for cells lacking mtDNA to receive the proliferation boost associated with Pde2p overexpression. (A) Overexpression of Tip41p, but not Pde2p, allows viability of cells lacking both Tom70p and mtDNA. Strains BY4741 (*WT*) and CDD897 (*tom70Δ*) were transformed with empty pRS426 vector, *2μ-PDE2* plasmid b89, or *2μ-TIP41* plasmid M489 and treated as in Fig. 8C. (B) Phosphorylation of S174 on Tom70p does not determine the outcome of mtDNA damage. Strain CDD913 (*tom70Δ*) was transformed with empty vector pRS314, plasmid b110 (pRS314-TOM70), plasmid b111 (pRS314-*tom70-S174Δ*), or plasmid b112 (pRS314-*tom70-S174E*). Resulting genotypes are shown. Strains were tested for their response to mtDNA deletion as in Fig. 1A except cells were incubated on solid SC-Trp medium for 2 d. (C) Tom70p is not upregulated in *ρ*^-^ cells overexpressing Pde2p. Whole cell extracts from strains CDD926 (*TOM70-GFP*) and CDD927 (*TOM70*) either overexpressing Pde2p from plasmid b89 or harboring an empty pRS426 vector and either containing or lacking mtDNA were analyzed by immunoblotting using antibodies recognizing GFP or hexokinase.

Next, we tested whether the petite-negative *tim18Δ* mutant, which lacks an IM component involved in the import of polytopic proteins into mitochondria [94], can remain viable after mtDNA loss upon PKA inhibition. Like *tom70Δ* mutants, *tim18Δ* mutants did not proliferate following mtDNA loss, even when carrying the *2μ-PDE2* plasmid (S4 Fig.). However, Pde2p overexpression also reduced the fitness of *tim18Δ* cells containing mtDNA, making this finding difficult to interpret. In the FY genetic background, in which *tim18Δ* mutants are more fit, overexpression of Pde2p did weakly suppress the petite-negative phenotype of *tim18Δ* cells (unpublished results), indicating that Tim18p is not totally required under all circumstances for *ρ*^-^ cells to receive the benefits of PKA inhibition.

Interestingly, PKA can phosphorylate proteins resident at mitochondria [95], including proteins at the TOM complex through which most proteins pass during import into these organelles [96–98]. Indeed, Tom70p is a PKA substrate. Since *tom70Δ ρ*^-^cells could not be rescued by PKA inhibition, we asked whether phosphorylation of a characterized PKA target site within Tom70p, S174, may be relevant to the increase in fitness provided to *ρ*^-^cells by Pde2p overproduction. *In vitro* import assays suggest that phosphorylation of Tom70p by PKA at S174 may inhibit Tom70p activity [98]. Therefore, if the benefit of PKA inhibition for *ρ*^-^ cells were due to reduced phosphorylation of Tom70p, then a S174A mutation that blocks PKA phosphorylation might increase the fitness of cells lacking mtDNA. However, Tom70p(S174A) expressed from a plasmid within *tom70Δ* mutant cells did not change the proliferation rate following mtDNA deletion (Fig. 9B). Moreover, the phosphomimetic S174E mutation also did not change the apparent division rate of cells lacking mtDNA. These results indicate that phosphorylation of residue S174 of Tom70p is not relevant to the effects of PKA reduction on *ρ*^-^ cells. These phosphomutant and phosphomimetic forms of Tom70p also provided no phenotypic effect at any temperature tested on either glucose-containing medium or on a non-fermentable medium (S5 Fig.).

We also tested the outcome of mutating PKA phosphorylation sites found upon two other TOM complex components: Tom22p and Tom40p. Specifically, T76 of Tom22p and S54 of Tom40p have been suggested to be phosphorylated in a PKA-dependent manner, with both phosphorylation events potentially inhibiting protein import [96,97]. However, the rate of proliferation following mtDNA loss was not altered upon expression of phosphomutant Tom22(T76A) and Tom40(S54A) proteins or upon the use of phosphomimetic Tom22(T76E) and Tom40(S54E) proteins (S6A Fig.). Moreover, alteration of these potential Tom22p and Tom40p phosphorylation sites did not notably alter proliferation of *ρ^+^* cells under any other condition tested (S6B Fig.).

Since Tom70p is apparently required for an increase in *ρ^-^* cell fitness due to PKA inhibition, we asked whether Tom70p might be upregulated at the transcriptional level in cells lacking mtDNA when Pde2p is overexpressed. However, *TOM70* transcription was nearly equivalent when comparing *ρ*^-^ cells overproducing Pde2p and *ρ*^-^ cells harboring an empty vector (S1 Table). Using a strain with the chromosomal *TOM70* allele tagged at the carboxyl-terminus with GFP, we asked whether the Tom70 polypeptide might be upregulated at the protein level upon PKA inhibition in *ρ*^-^ cells. The *TOM70-GFP* strain was increased in fitness upon Pde2p overexpression following mtDNA loss (S7 Fig.), indicating that the Tom70p-GFP is functional. However, Tom70p-GFP was not upregulated in *ρ*^-^ cells with elevated Pde2p levels (Fig. 9C). Therefore, Tom70p upregulation is not the mechanism by which PKA inhibition increase the fitness of cells lacking mtDNA.

As discussed above, Tom70p plays a prominent role in the mitochondrial import of polytopic proteins with internal targeting information, including Aac2p [99,100], which is required for proliferation following mtDNA loss [44,86,89]. Therefore, we asked whether levels of the Tom70p substrate Aac2p might be elevated upon Pde2p overexpression. However, *AAC2* did not appear significantly upregulated at the transcriptional level (S1 Table), and a functional, FLAG-tagged Aac2 protein was not increased in abundance in *ρ^-^* cells upon Pde2p overexpression (Fig. 10A). Therefore, an increase in Aac2p levels reliant upon by the Tom70 receptor seems unlikely to be the mechanism by which PKA benefits cells lacking mtDNA. However, we have not ruled out that a greater proportion of the total cellular pool of Aac2p is imported and properly assembled upon PKA inhibition in *ρ^-^* cells.

**Fig. 10:**
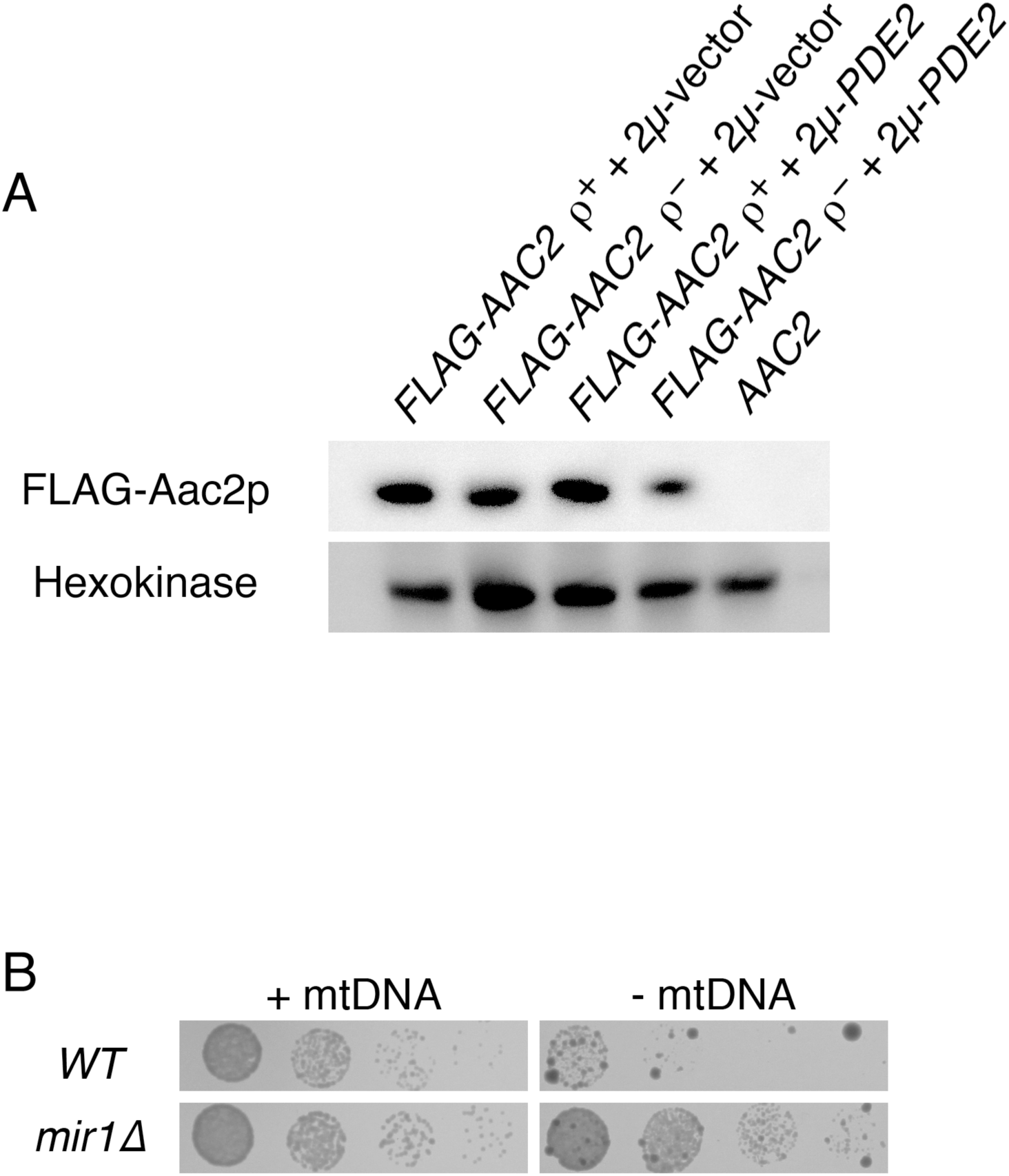
Pde2p overexpression does not benefit cells lacking mtDNA by Aac2p upregulation or through Mir1p upregulation. (A) Aac2p is not upregulated in *ρ*^-^ cells overproducing Pde2p. Whole cell extracts from *aac2Δ* strain CDD859 expressing FLAG-tagged Aac2p from plasmid b84 and either containing or deleted of mtDNA and either harboring 2*μ*-PDE2 plasmid b89 or empty vector pRS426 were analyzed using antibodies raised against the FLAG epitope tag or recognizing hexokinase. (B) Deletion of Mir1p provides an increase in proliferation rate to cells lacking mtDNA. Strains CDD862 (*WT*) and CDD863 (*mir1Δ*) were treated as in Fig. 1B, except *ρ^+^* cells were incubated for 1 d and *ρ*^-^ cells were incubated for 3 d.

As discussed above, ATP hydrolysis by the F_1_ sector of the ATP synthase is important for ΔΨ^mito^ generation [14,18,20,85–87], and ATP is moved to the matrix through the activity of Aac2p [44,86,89]. However, the electrogenic circuit required for ΔΨ^mito^ maintenance in cells lacking mtDNA is still incompletely understood, and little attention has been directed toward the inorganic phosphate generated by the F_1_-ATPase during ΔΨ^mito^ generation. Mir1p, the major phosphate transporter of the IM, is a substrate of Tom70p [101], and so we hypothesized that increased import of Mir1p might be a potential mechanism by which PKA reduction benefits *ρ^-^* cells. However, we found that *mir1Δ* cells lacking mtDNA proliferate more rapidly than *WT* cells lacking a mitochondrial genome (Fig. 10B), demonstrating that increased abundance or activity of Mir1p is not the mechanism by which mitochondrial dysfunction is reversed by reduced glucose sensation.

## Discussion

### Reduction of glucose sensation can benefit cells lacking mtDNA

Our results indicate that the sensation of glucose can control the outcome of mitochondrial dysfunction. Reduction of glucose concentration in the culture medium and inhibition of the major glucose-sensing signaling pathway of *S. cerevisiae* can increase the proliferation rate of cells lacking a mitochondrial genome. Previously published work supports our findings regarding glucose sensation: an increase in nuclear genomic instability prompted by mtDNA loss has been demonstrated to be suppressed by low glucose [102], and the same study reported that *mgr1Δ ρ^-^* colonies appear to be larger on glucose-restricted medium. Furthermore, a clone encoding only a portion of the adenylate cyclase Cyr1p, which is required for PKA activation, was identified as a potential dominant-negative suppressor of the inviability of *ρ*^-^ cells lacking the F_1_-ATPase (Prof. Xin Jie Chen, State University of New York Upstate Medical University, personal communication).

Since cells unable to perform OXPHOS rely on glycolysis for ATP generation, it may be surprising that a reduction in glucose sensation would increase the proliferation rate of *ρ*^-^ cells. Instead, one might expect that more glucose and glucose-dependent signaling would improve fitness under conditions of mitochondrial dysfunction. Supporting this idea, cancer cells defective in OXPHOS have been found to be badly affected by glucose limitation [103], and lactate production of mammalian *ρ*^-^ cells can be increased, suggesting increased glycolytic flux [20]. However, other evidence suggests that our findings could have general applicability to mitochondrial disease in metazoans. Specifically, glucose restriction appears to improve several parameters associated with mitochondrial dysfunction in a *Drosophila melanogaster* model of mitochondrial disease (Kemppainen et al., submitted). When considering the recent and exciting findings linking nutrition to the outcome of mitochondrial dysfunction across diverse experimental systems, further focus on dietary sugar and its effects on cells with damaged mitochondria may potentially lead to better outcomes for those afflicted by mitochondrial disease.

### What is the mechanism by which PKA inhibition increases fitness upon mtDNA loss?

We found that deletion of the Tpk3p isoform of PKA specifically provides an advantage to cells deleted of mtDNA, while Tpk1p and/or Tpk2p deletion was not beneficial for cells lacking mtDNA. Interestingly, Tpk3p was previously discovered to have a specific role among PKA isoforms in controlling mitochondrial function [104,105], although other PKA isoforms also impinge upon mitochondrial biogenesis [106]. Two comprehensive phosphoproteomic studies revealed nearly 200 proteins that may be direct or indirect targets of Tpk3p [34,107], and some of these may be found at mitochondria [95]. We note that Tpk3p inhibition likely provides only a portion of the benefits prompted by a reduction in the glucose-dependent signal, since *gpa2Δ ρ^-^* and *gpr1Δ ρ^-^* strains proliferate more quickly than *tpk3Δ ρ^-^* mutants. This result raises the possibility that other PKA isoforms also influence the outcome of mtDNA loss.

It is likely that the basis by which PKA inhibition promotes *ρ^-^* cell division differs from the mechanism by which perturbation of the Tap42p-controlled arm of the TORC1 pathway increases the proliferation of cells lacking a mitochondrial genome. Supporting this idea, overexpression of the Tap42p binding partner Tip41p suppresses the petite-negative phenotype of *tom70Δ ρ^-^* cell, while overexpression of Pde2p does not. Moreover, while overexpression of Pde2p leads to an increase in the expression of Hsp12p and Hsp26p, overexpression of Tip41p does not. However, even if the specific mechanisms by which *ρ*^-^ cells are benefited by these two pathways are initially divergent, is the final biochemical outcome at mitochondria also different? Our data do not provide a clear answer to this question. Deletion of *gpa2Δ* and *gpr1Δ,* while significantly increasing the proliferation rate of *ρ*^-^ cells, did not lead to increased mitochondrial localization of an *in vivo* reporter of mitochondrial protein import, while deletion of Tap42p-controlled phosphatases can lead to easily discernible re-localization of Cox4p(1-21)-GFP to *ρ^-^* mitochondria. The increase in mitochondrial protein import upon mutation of Tap42p-associated phosphatases is thought to be associated with an increase in the ΔΨ^mito^ [24], potentially indicating that the ΔΨ^mito^ of *ρ^-^* mitochondria is not significantly boosted by PKA inhibition. On the other hand, transcripts induced upon mtDNA loss, many of which are known to be determined by the magnitude of the ΔΨ^mito^ within *ρ^-^* mitochondria, are diminished in expression following mtDNA loss when Pde2p is overexpressed. Therefore, it remains possible that ΔΨ^mito^ is boosted by PKA inhibition in *ρ*^-^ cells, but not enough for Cox4p(1-21)-GFP to noticeably re-localize from the cytosol to mitochondria.

Moreover, while an increase in vacuolar pH is known to benefit *ρ^-^* cell proliferation through a potentially conserved mechanism, it is unlikely that reduced glucose sensation leads to improved fitness of cells lacking mtDNA by reducing endomembrane system acidity. First, overexpression of Pde2p did not cause decreased proliferation upon alkaline medium, a phenotype associated with a reduced ability to acidify the vacuole [79,80]. Moreover, a genome-wide analysis demonstrated no trend toward higher vacuolar pH upon PKA inhibition following deletion of Gpa2p, Gpr1p, or Tpk3p 108], and neither *gpa2Δ* nor *gpr1Δ* mutants phenocopy a mutant lacking the V_1_V_o_-ATPase with regard to Cox4p(1-21)-GFP localization in *ρ*^-^ cells.

Different genetic backgrounds of *S. cerevisiae* are characterized by divergent outcomes following mtDNA loss [109]. One gene that clearly plays a role in controlling the rate of proliferation following mtDNA loss is *MKT1.* The BY background, in which Pde2p overexpression provides more robust cell division, carries a derived allele of *MKT1* of potentially reduced function [110]. The W303 background, in which Pde2p overexpression does not promote *ρ^-^* proliferation, carries the ancestral allele of *MKT1* [109]. Since the ancestral allele of *MKT1* increases the fitness of cells lacking mtDNA, it may be that *MKT1* allele in the W303 background already protects cells from a specific pathological outcome of mtDNA damage, and therefore PKA inhibition by Pde2p overexpression can provide no further benefit. Alleles of other genes influencing mitochondrial biogenesis and function, such as *HAP1, SAL1,* and *CAT5,* also differ between the W303 background and the BY background [90,109,111] and may modify the response of *ρ*^-^ cells to PKA inhibition. Ongoing work in our laboratory is directed at mapping alleles that lead to background-specific effects on *ρ^-^* cell fitness.

We have ruled out numerous proteins and pathways as sole mediators of the effects of PKA inhibition on *ρ*^-^ cells, but clearly many other potential individual outcomes of PKA inhibition may remain to be tested. In addition, there may be a requirement that multiple PKA-controlled outcomes happen coincidentally in order for increased fitness of *ρ*^-^ cells to be achieved, yet further experimentation would be required in order to reveal this level of signaling complexity. Because mutants lacking Tom70p do not appear to respond positively to Pde2p overexpression following mtDNA loss, our attention is focused upon the abundance and function of substrates of the import pathway in which Tom70p participates.

### Derangement of amino acid synthesis and breakdown following mtDNA loss

The upregulation of the PDR, RTG, and IDR pathways are well-characterized outcomes of mtDNA damage in *S. cerevisiae* [24,36–39]. In this study, we found that numerous genes involved in arginine biosynthesis [42] are significantly upregulated upon mtDNA loss. Moreover, two genes that determine the abundance of aromatic amino acids [43], *ARO9* and *ARO10,* are repressed upon deletion of mtDNA. These changes were reversed by PKA inhibition. One caveat of our present approach to identifying transcriptional changes associated with mtDNA loss is that EtBr was present in the medium at the time *ρ*^-^ cells were harvested for RNA-seq analysis. However, our results are supported by a previous RNA-seq analysis of *ρ*^-^ cells never treated with EtBr [24], where several arginine biosynthesis enzymes were similarly activated and aromatic amino acid catabolism enzymes were also diminished. In that study, however, RNA-seq was not performed with replication, and therefore statistical significance for these changes was not previously observed.

These transcriptional changes hint at problems with amino acid homeostasis in *ρ*^-^ cells. Problems with maintaining amino acid levels may be a general consequence of mitochondrial dysfunction across many organisms, since synthesis of many amino acids depends upon carbon skeletons derived from the tricarboxylic acid cycle or requires other metabolites whose levels would be expected to change following a blockade of OXPHOS [4]. Previously, a genome-wide screen using *S. cerevisiae* revealed that mutations in the threonine biosynthesis pathway sensitize cells to mtDNA loss [15]. Very recently, cells with a dysfunctional respiratory chain were found to be deficient in aspartate [112,113]. Intriguingly, arginine levels are reported to be low in patients afflicted with Mitochondrial Encephalomyopathy, Lactic Acidosis, and Stroke-like episodes (MELAS), and some evidence might suggest that arginine supplementation can reduce MELAS symptoms [114]. While it may be very premature to connect our findings regarding transcriptional changes within the arginine biosynthesis pathway with human mitochondrial disease, particularly in light of differences in enzyme localization and regulation between yeast and man [115], amino acid level dysregulation upon mitochondrial DNA damage and its potential contribution to mitochondrial disease is clearly worthy of further attention.

## Materials and methods

### Yeast strains and culture conditions

YEPD medium contains 1% yeast extract, 2% bacteriological peptone, and 2% dextrose. YEPGE medium contains 1% yeast extract, 2% bacteriological peptone, 3% glycerol, and 3% ethanol. YEPLac medium contains 1% yeast extract, 2% bacteriological peptone, 1.2% NaOH, and a volume of lactic acid sufficient to bring the pH to 5.5. Synthetic complete (SC) medium contains 0.67% yeast nitrogen base without amino acids, 2% glucose, 0.1% casamino acids, 50 *μ*g/ml adenine hemisulfate, and either 25 *μ*g/ml uracil (SC-Trp) or 100 *μ*g/ml L-tryptophan (SC-Ura). We note that SC medium, replete with amino acids, was used to select for plasmid maintenance, since minimal medium can ameliorate the effects of mtDNA loss [93]. Solid media also contain 1.7% bacteriological agar. EtBr (Thermo Fisher Scientific) was used at a concentration of 25 *μ*g/ml, which is sufficient to destroy mtDNA [30]. However we use the “*ρ*^-^” rather than “*ρ^0^*” nomenclature throughout this work to describe cells lacking functional mtDNA, since these cultures were not further examined by DAPI or by other molecular assays to confirm total absence of mtDNA from all cells. Buffering of SC-Ura medium to pH 7.3 was accomplished using 100mM HEPES-KOH, pH 7.5 [75]; SC-Ura medium is typically pH 4.6. All cultures were performed at 30°C unless otherwise noted. For serial dilution assays, strains were maintained in logarithmic proliferation phase overnight in relevant media either lacking or containing EtBr, then diluted to an OD_600_ of 0.1. Four microliters of this dilution and three serial five-fold dilutions were spotted to solid medium. Figures are generated from plates photographed at incubation times that best demonstrate differences between control and experimental samples. Details of strain construction or acquisition are provided in S2 Table. Oligonucleotides used in this study are listed in S3 Table.

### Plasmid construction

All plasmid construction was performed by co-transformation of PCR products, along with *Not*I-cut vectors, into *S. cerevisiae* cells in order to allow homologous recombination between PCR product(s) and empty vector [116]. Plasmids were then recovered from yeast and amplified in bacterial strain DH5α. All plasmids used during the experiments found in this study, along with construction details, are described in S4 Table.

### Microscopy

Cells were visualized blind to genotype as in [16] to determine the number of cells in which mitochondrial localization of Cox4p(1-21)-GFP can be detected.

### Transcriptome analysis

BY4743 cells carrying plasmid pRS426 and maintaining mtDNA were cultured for at least three divisions in SC-Ura medium to ensure logarithmic phase proliferation. For cultures lacking mtDNA, BY4743 cells harboring plasmids pRS426, M489, or b89 were cultured in SC-Ura medium containing EtBr for 24 hours. For each sample type, two replicates were performed by different experimenters, for a total of eight biological samples.

Total RNA was extracted from 5 OD_600_ units of cells using the TRIzol reagent (Invitrogen) according to the manufacturer's instructions. RNA was treated with RNase-free DNase I (Thermo Fisher Scientific) at a concentration of 1 U/*μ*g to remove any remaining genomic DNA. mRNAs were purified from 2*μ*g total RNA using oligo (dT) magnetic beads and fragmented in fragmentation buffer using the TruSeq mRNA Sample Preparation Kit (Illumina). Fragmented RNA was used for first-strand cDNA synthesis using random hexameric primers, and the second strand was synthesized by using Second Strand Marking Mix (Illumina). Double-stranded cDNAs were purified with AMPure XP beads (Beckman Coulter) and eluted with re-suspension buffer, followed by adenine addition to the 3' end. Sequencing adaptors were ligated to the fragments, and cDNA fragments were enriched by PCR amplification. Enriched cDNA libraries were used for cluster generation and sequencing. Paired-end sequencing of the cDNA libraries was performed using the Illumina Miseq platform (Illumina). All sequence data are paired-end, 2 × 75bp. Image processing, base calling, and quality value calculation were performed by the Illumina data processing pipeline (v1.5). High quality reads were saved in FASTQ format.

Bioinformatic analysis of sequencing reads was performed using the resources of the Galaxy Project [117]. During our analysis, fastq_groomer on FASTQ files (Galaxy Tool Version 1.0.4) was performed on FASTQ data files, followed by the use of TopHat 2.0.14 [118,119] on groomed FASTQ files (Galaxy tool version 0.9) with mean inner distance between mate pairs set to 300 bases and standard deviation for the distance between mate pairs set to 50 bases, reporting of discordant pair alignments, and the use of *S. cerevisiae* reference genome R64-2-1 [120]. Moreover, preset TopHat settings were toggled, read group was not specified, and job resource parameters were set to 'no.' Results of the TopHat analysis were then analyzed within Cuffdiff 2.2.1.1 (Galaxy Tool Version 2.2.1.1) with transcript annotation provided by the GFF file for *S. cerevisiae* reference genome R64-2-1 [121], geometric library normalization, per-condition dispersion estimation methods, a false discovery rate of 0.05, and a minimum alignment count of 10. Multi-read correct was utilized, as was bias correction using the R64-2-1 reference sequence. Read group datasets was not toggled, count-based output files were included, and Cufflinks effective length correction was applied. No additional parameters were set. Formatted data are found in S1 Table, and raw Cufflinks output and sequence reads can be found under GEO accession number GSE71252.

### Immunoblotting

For cultures containing mtDNA, cells were grown overnight in SC-Ura to ensure logarithmic phase of proliferation. For cultures lacking mtDNA, cells were incubated for 24 hr at logarithmic phase in medium containing EtBr. Protein extraction was performed essentially as described in [122]. Briefly, 5 mL of cells in culture were boiled for 3 min, and the cell pellet was resuspended in 200 *μ*L 1 X TE buffer (10mM Tris-HCl, pH 8.0; 1 mM EDTA), followed by 200 *μ*L of 0.2 N NaOH. The sample was incubated at room temperature for 5 min, followed by centrifugation and addition of 1 X SDS-PAGE sample buffer to the pellet. Equal OD units of cells were loaded onto acrylamide gels and proteins were separated using a Bis-Tris-based gel system [123] with 1 X MOPS-SDS running buffer (National Diagnostics) before semi-dry transfer onto a PVDF membrane (Thermo Fisher Scientific). Equivalent protein loading and transfer was confirmed using the MemCode Reversible Protein Stain Kit (Pierce), and blocking was performed with 0.5% milk blocking buffer solution in TBST (10mM Tris-HCl pH 7.5, 150 mM NaCl, 0.05% Tween 20). Primary antibodies were added in 0.1% milk blocking buffer solution in TBST. Mouse anti-FLAG M2 antibody (Sigma; F1804) was used at a dilution of 1:10000 to detect FLAG-tagged Aac2p. Rabbit anti-GFP antibody (Cell Signaling Technology; 2956) was used at 1:5000 to detect Tom70p-GFP. Rabbit anti-hexokinase antibody was used at 1:20000 to detect hexokinase (a gift of Rob Jensen; JH2616). Secondary antibodies (Cell Signaling Technology; 7074 and 7076) were used at 1:10000 to allow subsequent chemiluminescent detection of relevant proteins.

**Figure legends**

**S1 Fig.:**
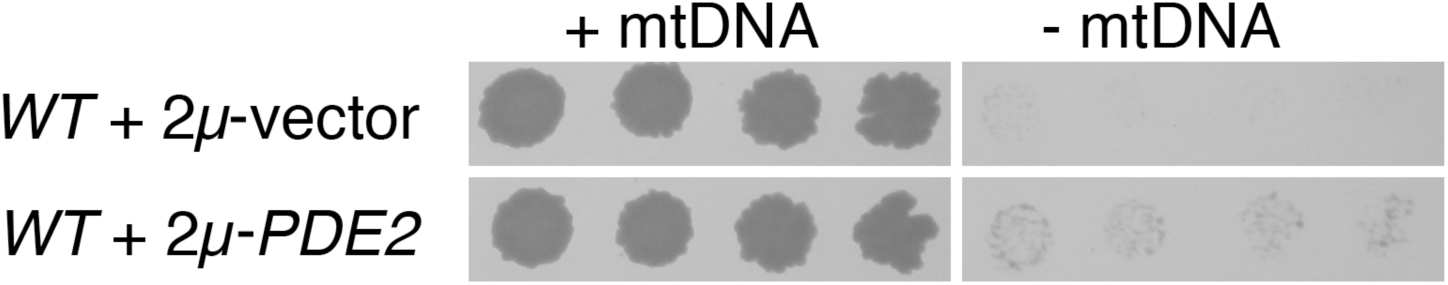
Cells cultured overnight in EtBr lose functional mtDNA. Spot-dilution tests from Fig. 1A were replica-plated to YEPGE medium and incubated for 2 d.

**S2 Fig.:**
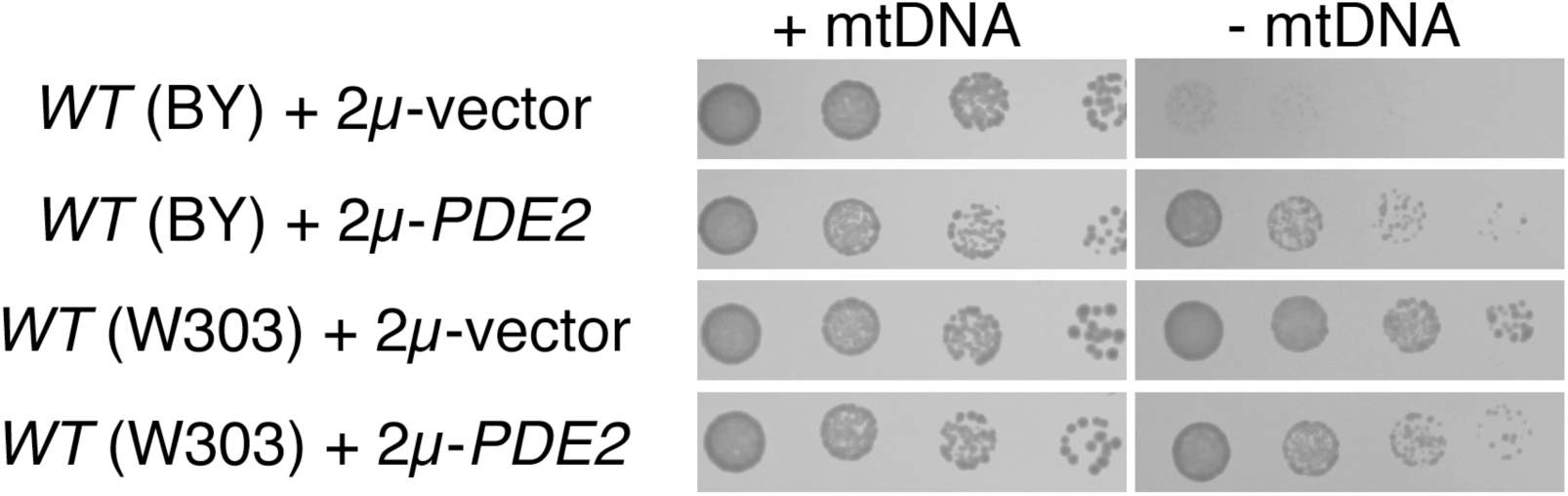
The benefits provided to cells deleted of the mitochondrial genome by Pde2p overexpression are dependent upon yeast genetic background. Strains BY4741 (*WT*, BY background) and BMA64-1A (*WT*, W303 background) were treated as in Fig. 1A.

**S3 Fig.:**
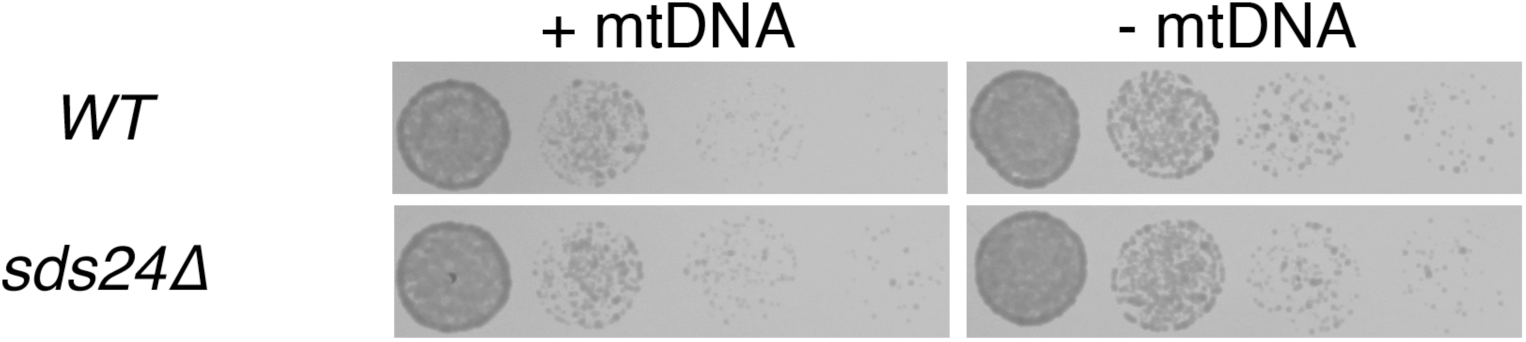
Mutants lacking Sds24p do not exhibit a proliferation defect after mtDNA loss. Strains CDD463 (*WT*) and CDD912 (*sds24Δ*) were treated as in Fig. 1B, except *ρ^+^* cells were incubated on solid YEPD medium for 1 d, and *ρ*^-^ cells were incubated for 2 d.

**S4 Fig.:**
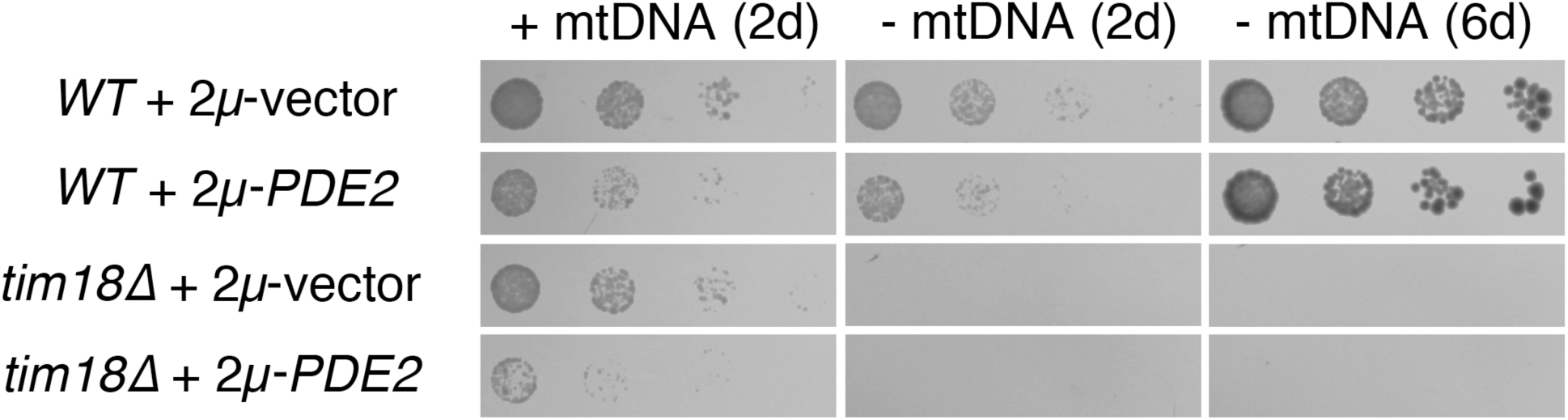
Cells of the BY background lacking Tim18p are unresponsive to the benefits provided by Pde2p overexpression to *ρ*^-^cells. Strains BY4741 (*WT*) and CDD9 (*tim18Δ*) were treated as in Fig. 8C, except that cultures were proliferated and maintained at 37°C to circumvent the cold-sensitivity of the *tim18Δ* mutant.

**S5 Fig.:**
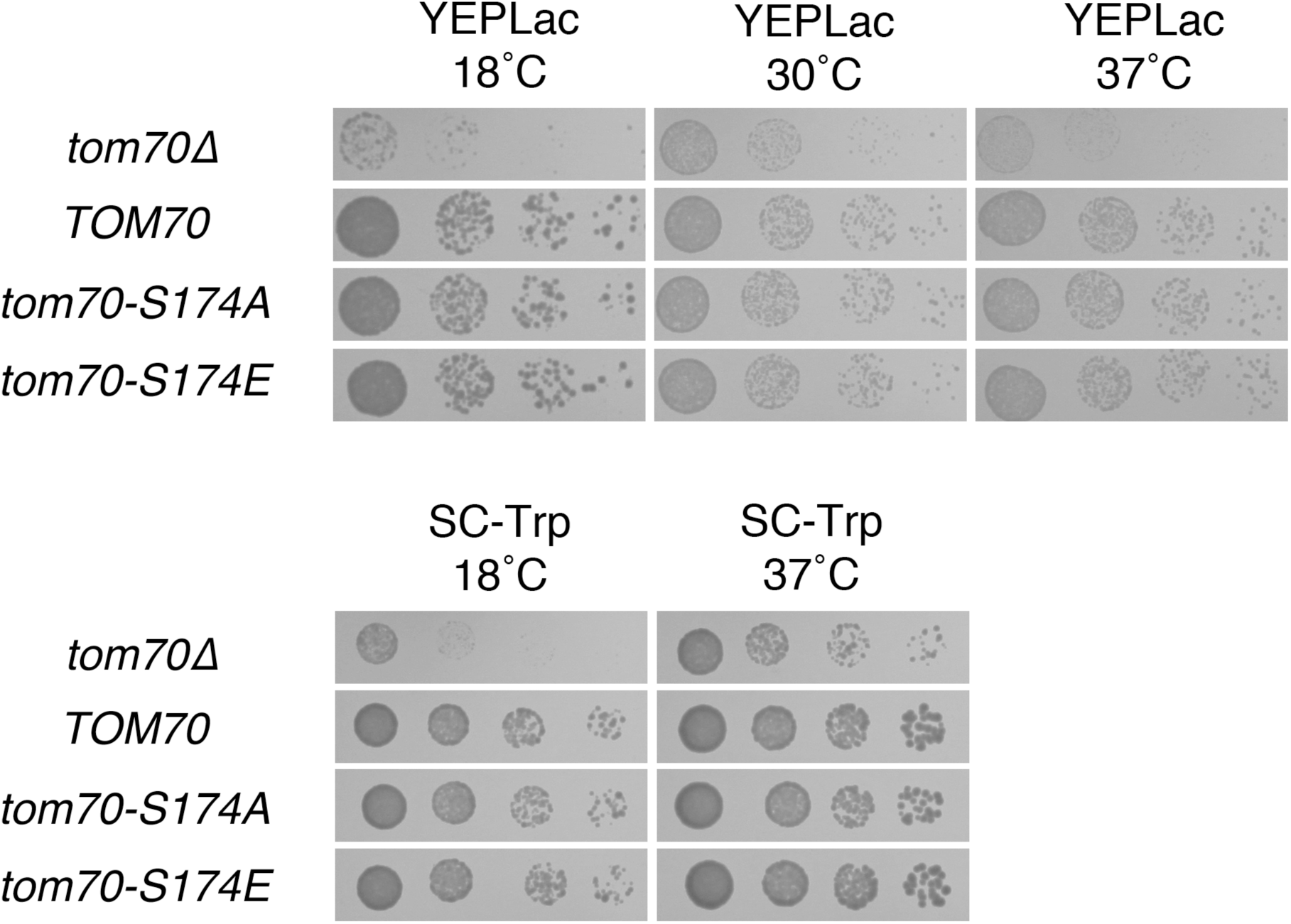
Phosphorylation of S174 of Tom70p has no apparent consequence for cellular proliferation. *ρ^+^* cells transformants used in Fig. 9 were also plated to the media indicated and incubated for 2 d (SC-Trp at 37°C, YEPLac at 30°C, YEPLac at 37°C); 4 d (SD-Trp at 18°C); or 7 d (YEPLac at 18°C).

**S6 Fig.:**
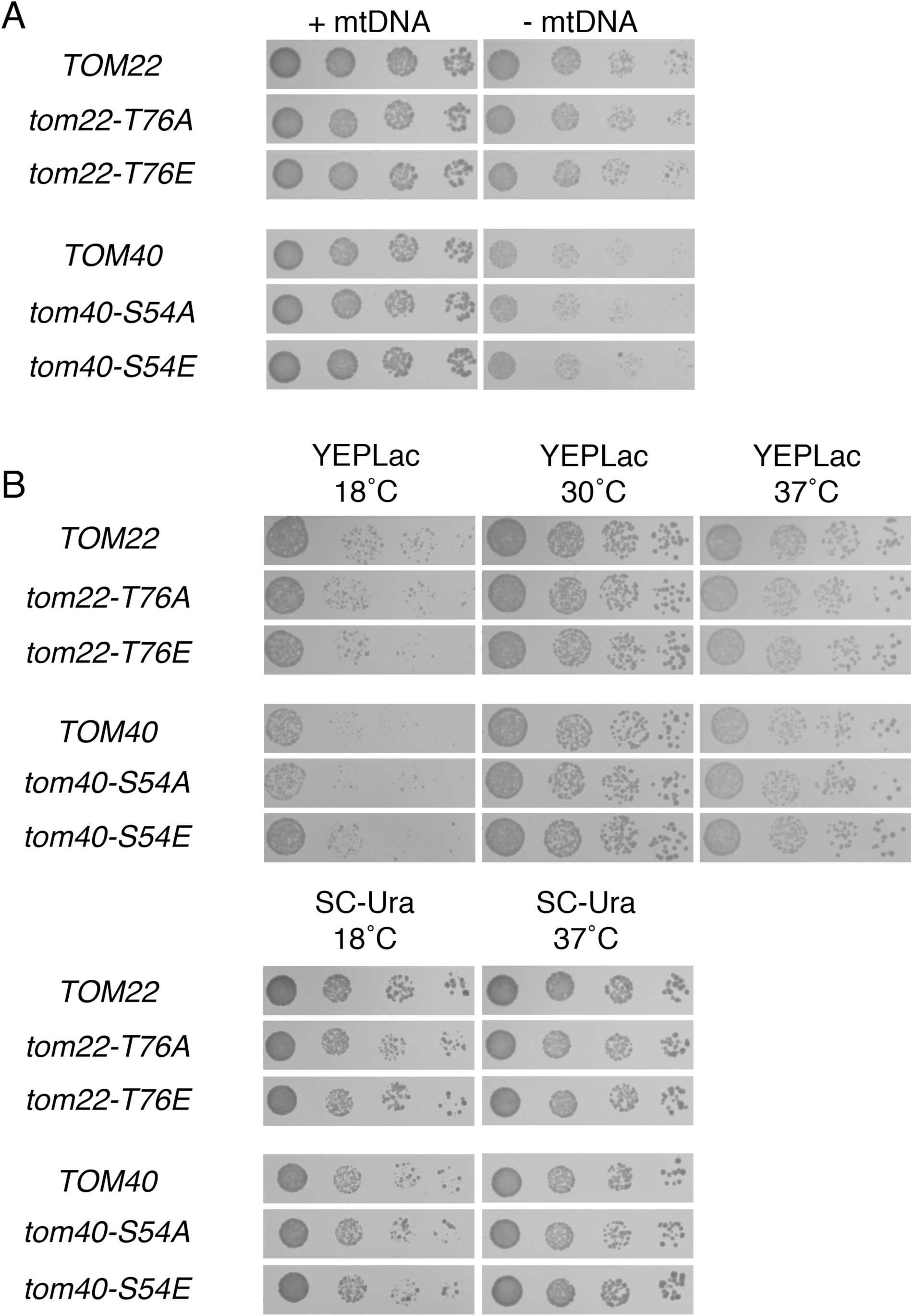
Known PKA phosphorylation target sites on Tom40p and Tom22p are not relevant for proliferation rate following mtDNA loss or under several other culture conditions. (A) Phosphorylation of T76 on Tom22p and phosphorylation of S54 of Tom40p do not determine the outcome of mtDNA damage. Strains CDD866 (*TOM22*), CDD867 (*tom22-T76A*), CDD868 (*tom22-T76E*), CDD869 (*TOM40*), CDD870 (*tom40-S54A*), and CDD871 (*tom40-S54E*), each manifesting chromosomal deletions complemented by plasmid-borne variants, were treated as in Fig. 1A. Relevant genotypes are shown. (B) Phosphorylation of T76 on Tom22p and of S54 of Tom40p have no apparent consequence for cellular proliferation. *ρ^+^* cultures used in (A) were plated to the media indicated and incubated for 2 d (SC-Ura at 37°C); 3 d (YEPLac at 30°C, YEPLac at 37°C), or 5 d (SC-Ura at 18°C, YEPLac at 18°C).

**S7 Fig.:**
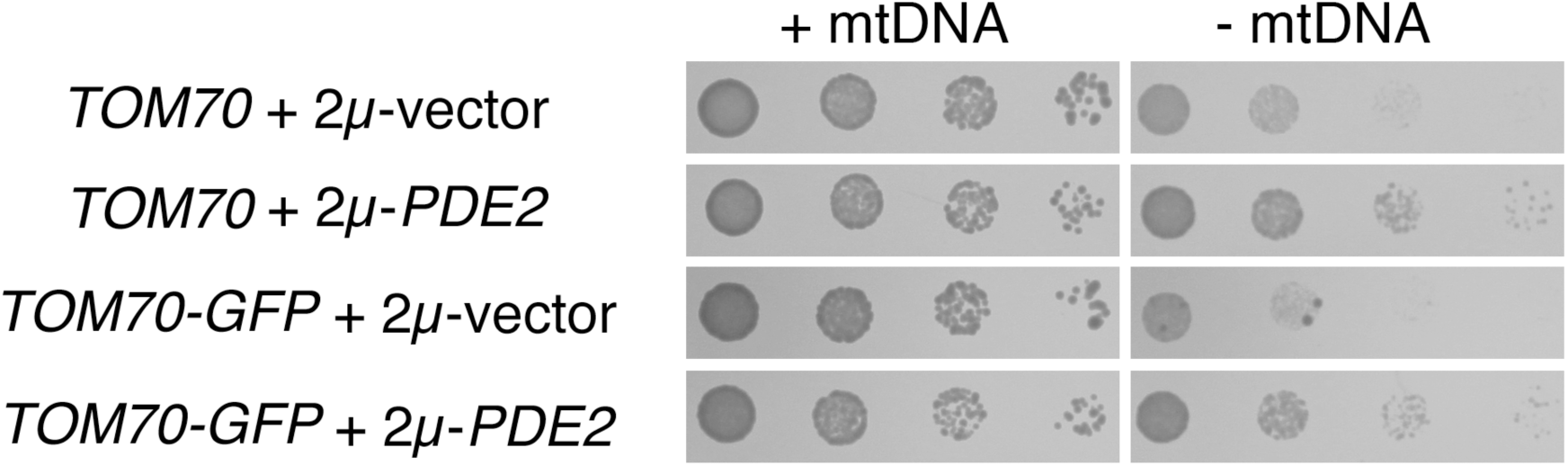
A strain expressing GFP-tagged Tom70p and lacking mtDNA responds to PKA inhibition. Strains CDD926 (*TOM70-GFP*) and CDD927 (*TOM70*) were treated as in Fig. 1A.

**S1 Tab.:** RNA-Seq data.

**S2 Tab.:** Strains used in this study.

**S3 Tab.:** Oligonucleotides used in this study.

**S4 Tab.:** Plasmids used within experiments in this study.

## Acknowledgements

We thank Gülayşe ince Dunn, Abdurrahman Keskin, Arwa Bayrakdar, and Funda Kar for comments on this manuscript, and we appreciate Xin Jie Chen's (State University of New York Upstate Medical School) communication of unpublished results. We thank Rob Jensen (Johns Hopkins School of Medicine) for providing antibodies to hexokinase and Maya Schuldiner (Weizmann Institute) for *hsf1-DAmP and TOM70-GFP* strains. This work was supported by a European Research Council Starting Grant (637649-RevMito) to CDD, by a European Molecular Biology Organization Installation Grant to CDD, by an Istanbul Development Agency grant (ISTKA-TR/14/EVK/0039) to IHK, and by Koç University.

## Author contributions

Conceived and designed the experiments: EA CDD. Performed the experiments: EA MT CDD. Analyzed the data: EA MT IHK CDD. Contributed reagents/materials/analysis tools: EA GG GB IHK CDD. Wrote the paper: EA MT CDD.

## References

1. Kispal G, Sipos K, Lange H, Fekete Z, Bedekovics T, Janaky T, et al. Biogenesis of cytosolic ribosomes requires the essential iron-sulphur protein Rli1p and mitochondria. EMBO J. 2005 ed. 2005;24: 589–598. doi:10.1038/sj.emboj.7600541

2. Lill R, Hoffmann B, Molik S, Pierik AJ, Rietzschel N, Stehling O, et al. The role of mitochondria in cellular iron–sulfur protein biogenesis and iron metabolism. BBA-Molecular Cell Research. Elsevier B.V; 2012;: 1–18. doi:10.1016/j.bbamcr.2012.05.009

3. Gray MW. The Pre-Endosymbiont Hypothesis: A New Perspective on the Origin and Evolution of Mitochondria. Cold Spring Harbor Perspectives in Biology. 2014;6: a016097–a016097. doi:10.1101/cshperspect.a016097

4. Voet D, Voet JG. Biochemistry, 4th Edition. 4 ed. Wiley Global Education; 2010.

5. Schaefer AM, McFarland R, Blakely EL, He L, Whittaker RG, Taylor RW, et al. Prevalence of mitochondrial DNA disease in adults. Ann Neurol. 2007 ed. 2008;63: 35–39. doi:10.1002/ana.21217

6. Payne BAI, Wilson IJ, Hateley CA, Horvath R, Santibanez-Koref M, Samuels DC, et al. Mitochondrial aging is accelerated by anti-retroviral therapy through the clonal expansion of mtDNA mutations. Nat Genet. 2011 ed. 2011; 43: 1–7. doi:10.1038/ng.863

7. Lebrecht D, Walker UA. Role of mtDNA lesions in anthracycline cardiotoxicity. Cardiovascular toxicology. 2007;7: 108–113. doi:10.1007/s12012-007-0009-1

8. Carew JS, Zhou Y, Albitar M, Carew JD, Keating MJ, Huang P. Mitochondrial DNA mutations in primary leukemia cells after chemotherapy: clinical significance and therapeutic implications. Leukemia. 2003;17: 1437–1447. doi:10.1038/sj.leu.2403043

9. Payne BAI, Chinnery PF. Mitochondrial dysfunction in aging: Much progress but many unresolved questions. Biochim Biophys Acta. 2015. doi:10.1016/j.bbabio.2015.05.022

10. Ruetenik A, Barrientos A. Dietary restriction, mitochondrial function and aging: from yeast to humans. Biochim Biophys Acta. 2015. doi:10.1016/j.bbabio.2015.05.005

11. Pfeffer G, Horvath R, Klopstock T, Mootha VK, Suomalainen A, Koene S, et al. New treatments for mitochondrial disease-no time to drop our standards. Nat Rev Neurol. 2013;9: 474–481. doi:10.1038/nrneurol.2013.129

12. Suomalainen A. Seminars in Fetal & Neonatal Medicine. Seminars in Fetal and Neonatal Medicine. Elsevier Ltd; 2011; 16: 236–240. doi:10.1016/j.siny.2011.05.003

13. Bulder CJ. Lethality of the petite mutation in petite negative yeasts. Antonie Van Leeuwenhoek. 1964;30: 442–454.

14. Chen XJ, Clark-Walker GD. The petite mutation in yeasts: 50 years on. Int Rev Cytol. 2000; 194: 197–238.

15. Dunn CD, Lee MS, Spencer FA, Jensen RE. A genomewide screen for petite-negative yeast strains yields a new subunit of the i-AAA protease complex. Mol Biol Cell. 2005 ed. 2006;17: 213–226. doi:10.1091/mbc.E05-06-0585

16. Garipler G, Dunn CD. Defects associated with mitochondrial DNA damage can be mitigated by increased vacuolar pH in *Saccharomyces cerevisiae*. Genetics. 2013;194: 285–290. doi:10.1534/genetics. 113.149708

17. Baker N, Hamilton G, Wilkes JM, Hutchinson S, Barrett MP, Horn D. Vacuolar ATPase depletion affects mitochondrial ATPase function, kinetoplast dependency, and drug sensitivity in trypanosomes. Proc Natl Acad Sci USA. 2015;112: 9112–9117. doi:10.1073/pnas.1505411112

18. Giraud MF, Velours J. The absence of the mitochondrial ATP synthase delta subunit promotes a slow growth phenotype of rho-yeast cells by a lack of assembly of the catalytic sector F1. Eur J Biochem. Blackwell Science Ltd; 1997;245: 813–818. doi:10.1111/j.1432-1033.1997.00813.x

19. Dean S, Gould MK, Dewar CE, Schnaufer AC. Single point mutations in ATP synthase compensate for mitochondrial genome loss in trypanosomes. Proceedings of the National Academy of Sciences. 2013;110: 14741–14746. doi:10.1073/pnas. 1305404110

20. Buchet K, Godinot C. Functional F1-ATPase essential in maintaining growth and membrane potential of human mitochondrial DNA-depleted rho degrees cells. J Biol Chem. 1998;273: 22983–22989.

21. Cunningham JT, Rodgers JT, Arlow DH, Vazquez F, Mootha VK, Puigserver P. mTOR controls mitochondrial oxidative function through a YY1-PGC-1alpha transcriptional complex. Nature. 2007;450: 736–740. doi:10.1038/nature06322

22. Zong H, Ren JM, Young LH, Pypaert M, Mu J, Birnbaum MJ, et al. AMP kinase is required for mitochondrial biogenesis in skeletal muscle in response to chronic energy deprivation. Proc Natl Acad Sci USA. 2002;99: 15983–15987. doi:10.1073/pnas.252625599

23. Hedbacker K, Carlson M. SNF1/AMPK pathways in yeast. Front Biosci. 2008;13: 2408–2420.

24. Garipler G, Mutlu N, Lack NA, Dunn CD. Deletion of conserved protein phosphatases reverses defects associated with mitochondrial DNA damage in *Saccharomyces cerevisiae*. Proceedings of the National Academy of Sciences. 2014;111: 1473–1478. doi:10.1073/pnas.1312399111

25. Peng M, Ostrovsky J, Kwon YJ, Polyak E, Licata J, Tsukikawa M, et al. Inhibiting cytosolic translation and autophagy improves health in mitochondrial disease. Human Molecular Genetics. 2015. doi:10.1093/hmg/ddv207

26. Tain LS, Mortiboys H, Tao RN, Ziviani E, Bandmann O, Whitworth AJ. Rapamycin activation of 4E-BP prevents parkinsonian dopaminergic neuron loss. Nat Neurosci. 2009;12: 1129–1135. doi:10.1038/nn.2372

27. Johnson SC, Yanos ME, Kayser EB, Quintana A, Sangesland M, Castanza A, et al. mTOR Inhibition Alleviates Mitochondrial Disease in a Mouse Model of Leigh Syndrome. Science. 2013;342: 1524–1528. doi:10.1126/science.1244360

28. Broach JR. Nutritional Control of Growth and Development in Yeast. 2012;192: 73–105. doi:10.1534/genetics. 111.135731

29. Sass P, Field J, Nikawa J, Toda T, Wigler M. Cloning and characterization of the high-affinity cAMP phosphodiesterase of *Saccharomyces cerevisiae*. Proc Natl Acad Sci USA. National Academy of Sciences; 1986;83: 9303–9307.

30. Goldring ES, Grossman LI, Krupnick D, Cryer DR, Marmur J. The petite mutation in yeast. Loss of mitochondrial deoxyribonucleic acid during induction of petites with ethidium bromide. J Mol Biol. 1970;52: 323–335. doi:10.1016/0022-2836(70)90033-1

31. Brachmann CB, Davies A, Cost GJ, Caputo E, Li J, Hieter P, et al. Designer deletion strains derived from Saccharomyces cerevisiae S288C: a useful set of strains and plasmids for PCR-mediated gene disruption and other applications. Yeast. 1998;14: 115–132.

32. Thomas BJ, Rothstein R. Elevated recombination rates in transcriptionally active DNA. Cell. 1989;56: 619–630.

33. Garrett S, Broach J. Loss of Ras activity in Saccharomyces cerevisiae is suppressed by disruptions of a new kinase gene, YAKI, whose product may act downstream of the cAMP-dependent protein kinase. Genes Dev. 1989;3: 1336–1348.

34. Ptacek J, Devgan G, Michaud G, Zhu H, Zhu X, Fasolo J, et al. Global analysis of protein phosphorylation in yeast. Nature. 2005;438: 679–684. doi:10.1038/nature04187

35. Xue Y, Batlle M, Hirsch JP. GPR1 encodes a putative G protein-coupled receptor that associates with the Gpa2p Galpha subunit and functions in a Ras-independent pathway. EMBO J. 1998;17: 1996–2007. doi:10.1093/emboj/17.7.1996

36. Veatch JR, Mcmurray MA, Nelson ZW, Gottschling DE. Mitochondrial dysfunction leads to nuclear genome instability via an iron-sulfur cluster defect. Cell. 2009;137: 1247–1258. doi:10.1016/j.cell.2009.04.014

37. Hallstrom TC. Multiple Signals from Dysfunctional Mitochondria Activate the Pleiotropic Drug Resistance Pathway in *Saccharomyces cerevisiae*. 2000;275: 37347–37356. doi:10.1074/jbc.M007338200

38. Epstein CB, Waddle JA, Hale W, Davé V, Thornton J, Macatee TL, et al. Genome-wide responses to mitochondrial dysfunction. Mol Biol Cell. American Society for Cell Biology; 2001; 12: 297–308.

39. Liao X, Butow RA. RTG1 and RTG2: two yeast genes required for a novel path of communication from mitochondria to the nucleus. Cell. 1993;72: 61–71.

40. Miceli MV, Jiang JC, Tiwari A, Rodriguez-Quiñones JF, Jazwinski SM. Loss of mitochondrial membrane potential triggers the retrograde response extending yeast replicative lifespan. Front Genet. 2011; 2: 102. doi:10.3389/fgene.2011.00102

41. Di Como CJ, Arndt KT. Nutrients, via the Tor proteins, stimulate the association of Tap42 with type 2A phosphatases. Genes Dev. Cold Spring Harbor Lab; 1996;10: 1904–1916. doi:10.1101/gad.10.15.1904

42. Ljungdahl PO, Daignan-Fornier B. Regulation of amino acid, nucleotide, and phosphate metabolism in *Saccharomyces cerevisiae*. Genetics. Genetics Society of America; 2012;190: 885–929. doi:10.1534/genetics.111.133306

43. Lee K, Hahn J-S. Interplay of Aro80 and GATA activators in regulation of genes for catabolism of aromatic amino acids in *Saccharomyces cerevisiae*. Mol Microbiol. 2013;88: 1120–1134. doi:10.1111/mmi.12246

44. Dupont CH, Mazat JP, Guerin B. The role of adenine nucleotide translocation in the energization of the inner membrane of mitochondria isolated from rho + and rho degree strains of Saccharomyces cerevisiae. Biochemical and Biophysical Research Communications. 1985;132: 1116–1123.

45. Reid GA, Schatz G. Import of proteins into mitochondria. Yeast cells grown in the presence of carbonyl cyanide m-chlorophenylhydrazone accumulate massive amounts of some mitochondrial precursor polypeptides. J Biol Chem. 1982;257: 13056–13061.

46. Pedruzzi I, Burckert N, Egger P, De Virgilio C. *Saccharomyces cerevisiae* Ras/cAMP pathway controls post-diauxic shift element-dependent transcription through the zinc finger protein Gis1. EMBO J. EMBO Press; 2000;19: 2569–2579. doi:10.1093/emboj/19.11.2569

47. Görner W, Durchschlag E, Wolf J, Brown EL, Ammerer G, Ruis H, et al. Acute glucose starvation activates the nuclear localization signal of a stress-specific yeast transcription factor. EMBO J. EMBO Press; 2002;21: 135–144. doi:10.1093/emboj/21.1.135

48. Görner W, Durchschlag E, Martinez-Pastor MT, Estruch F, Ammerer G, Hamilton B, et al. Nuclear localization of the C2H2 zinc finger protein Msn2p is regulated by stress and protein kinase A activity. Genes Dev. Cold Spring Harbor Laboratory Press; 1998;12: 586–597.

49. Swinnen E, Wanke V, Roosen J, Smets B, Dubouloz F, Pedruzzi I, et al. Rim15 and the crossroads of nutrient signalling pathways in *Saccharomyces cerevisiae*. Cell Div. BioMed Central Ltd; 2006;1: 3. doi:10.1186/1747-1028-1-3

50. Morano KA, Grant CM, Moye-Rowley WS. The Response to Heat Shock and Oxidative Stress in *Saccharomyces cerevisiae*. Genetics. 2012;190: 1157–1195. doi:10.1534/genetics. 111.128033

51. Ferguson SB, Anderson ES, Harshaw RB, Thate T, Craig NL, Nelson HC. Protein kinase A regulates constitutive expression of small heat-shock genes in an Msn2/4p-independent and Hsf1p-dependent manner in *Saccharomyces cerevisiae*. Genetics. 2005;169: 1203–1214. doi:10.1534/genetics.104.034256

52. Breslow DK, Cameron DM, Collins SR, Schuldiner M, Stewart-Ornstein J, Newman HW, et al. A comprehensive strategy enabling high-resolution functional analysis of the yeast genome. Nat Meth. 2008;5: 711–718. doi:10.1038/nmeth. 1234

53. Yamamoto A, Mizukami Y, Sakurai H. Identification of a novel class of target genes and a novel type of binding sequence of heat shock transcription factor in *Saccharomyces cerevisiae*. J Biol Chem. American Society for Biochemistry and Molecular Biology; 2005;280: 11911–11919. doi:10.1074/jbc.M411256200

54. Reinders A, Burckert N, Boller T, Wiemken A, De Virgilio C. Saccharomyces cerevisiae cAMP-dependent protein kinase controls entry into stationary phase through the Rim15p protein kinase. Genes Dev. 1998;12: 2943–2955.

55. Lee P, Kim MS, Paik S-M, Choi S-H, Cho B-R, Hahn J-S. Rim15-dependent activation of Hsf1 and Msn2/4 transcription factors by direct phosphorylation in *Saccharomyces cerevisiae*. FEBS Lett. Federation of European Biochemical Societies; 2013;587: 3648–3655. doi:10.1016/j.febslet.2013.10.004

56. Amoros M, Estruch F. Hsf1p and Msn2/4p cooperate in the expression of Saccharomyces cerevisiae genes HSP26 and HSP104 in a gene- and stress type-dependent manner. Mol Microbiol. 2001;39: 1523–1532.

57. Lippman SI, Broach JR. Protein kinase A and TORC1 activate genes for ribosomal biogenesis by inactivating repressors encoded by Dot6 and its homolog Tod6. Proceedings of the National Academy of Sciences. 2009;106: 19928–19933. doi:10.1073/pnas.0907027106

58. Huber A, French SL, Tekotte H, Yerlikaya S, Stahl M, Perepelkina MP, et al. Sch9 regulates ribosome biogenesis via Stb3, Dot6 and Tod6 and the histone deacetylase complex RPD3L. EMBO J. 2011; 30: 3052–3064. doi:10.1038/emboj.2011.221

59. Deminoff SJ, Howard SC, Hester A, Warner S, Herman PK. Using substrate-binding variants of the cAMP-dependent protein kinase to identify novel targets and a kinase domain important for substrate interactions in *Saccharomyces cerevisiae*. Genetics. 2006;173: 1909–1917. doi:10.1534/genetics.106.059238

60. Wang X, Chen XJ. A cytosolic network suppressing mitochondria-mediated proteostatic stress and cell death. Nature. 2015. doi:10.1038/nature14859

61. Wang X, Zuo X, Kucejova B, Chen XJ. Reduced cytosolic protein synthesis suppresses mitochondrial degeneration. Nat Cell Biol. 2008;10: 1090–1097. Available: http://eutils.ncbi.nlm.nih.gov/entrez/eutils/elink.fcgi?dbfrom=pubmed&id=19160490&retmode=ref&cmd=prlinks

62. Maleszka R, Clark-Walker GD. A petite positive strain of Kluyveromyces lactis has a 300 kb deletion in the rDNA cluster. Curr Genet. 1989;16: 429–432.

63. Pluta K, Lefebvre O, Martin NC, Smagowicz WJ, Stanford DR, Ellis SR, et al. Maf1p, a negative effector of RNA polymerase III in *Saccharomyces cerevisiae*. Mol Cell Biol. American Society for Microbiology; 2001; 21: 5031–5040. doi:10.1128/MCB.21. 15.5031-5040.2001

64. Upadhya R, Lee J, Willis IM. Maf1 is an essential mediator of diverse signals that repress RNA polymerase III transcription. Molecular Cell. 2002;10: 1489–1494.

65. Moir RD, Lee J, Haeusler RA, Desai N, Engelke DR, Willis IM. Protein kinase A regulates RNA polymerase III transcription through the nuclear localization of Maf1. Proc Natl Acad Sci USA. National Acad Sciences; 2006;103: 15044–15049. doi:10.1073/pnas.0607129103

66. Hedbacker K, Townley R, Carlson M. Cyclic AMP-dependent protein kinase regulates the subcellular localization of Snf1-Sip1 protein kinase. Mol Cell Biol. 2004;24: 1836–1843. doi:10.1128/MCB.24.5.1836–1843.2004

67. Yakura M, Ishikura Y, Adachi Y, Kawamukai M. Involvement of Moc1 in sexual development and survival of Schizosaccharomyces pombe. Bioscience, Biotechnology, and Biochemistry. Japan Society for Bioscience, Biotechnology, and Agrochemistry; 2006;70: 1740–1749. doi:10.1271/bbb.60088

68. Anaul Kabir M, Kaminska J, Segel GB, Bethlendy G, Lin P, Seta Della F, et al. Physiological effects of unassembled chaperonin Cct subunits in the yeast *Saccharomyces cerevisiae*. Yeast. 2005;22: 219–239. doi:10.1002/yea.1210

69. Hanyu Y, Imai KK, Kawasaki Y, Nakamura T, Nakaseko Y, Nagao K, et al. Schizosaccharomyces pombe cell division cycle under limited glucose requires Ssp1 kinase, the putative CaMKK, and Sds23, a PP2A-related phosphatase inhibitor. Genes Cells. 2009 ed. 2009;14: 539–554. doi:10.1111/j.1365-2443.2009.01290.x

70. Teng X, Dayhoff-Brannigan M, Cheng W-C, Gilbert CE, Sing CN, Diny NL, et al. Genome-wide consequences of deleting any single gene. Mol Cell. 2013;52: 485–494. doi:10.1016/j.molcel.2013.09.026

71. Mitchell SF, Parker R. Principles and Properties of Eukaryotic mRNPs. Molecular Cell. Elsevier Inc; 2014;54: 547–558. doi:10.1016/j.molcel.2014.04.033

72. Tudisca V, Recouvreux V, Moreno S, Boy-Marcotte E, Jacquet M, Portela P. Differential localization to cytoplasm, nucleus or P-bodies of yeast PKA subunits under different growth conditions. Eur J Cell Biol. 2010;89: 339–348. doi:10.1016/j.ejcb.2009.08.005

73. Ramachandran V, Shah KH, Herman PK. The cAMP-Dependent Protein Kinase Signaling Pathway Is a Key Regulator of P Body Foci Formation. Molecular Cell. Elsevier Inc; 2011;43: 973–981. doi:10.1016/j.molcel.2011.06.032

74. Teixeira D, Parker R. Analysis of P-body assembly in *Saccharomyces cerevisiae*. Mol Biol Cell. American Society for Cell Biology; 2007;18: 2274–2287. doi:10.1091 /mbc.E07-03-0199

75. Hughes AL, Gottschling DE. An early age increase in vacuolar pH limits mitochondrial function and lifespan in yeast. Nature. 2012;492: 261–265. doi:10.1038/nature11654

76. Elbaz-Alon Y, Rosenfeld-Gur E, Shinder V, Futerman AH, Geiger T, Schuldiner M. Short Article. Dev Cell. Elsevier Inc; 2014;30: 95–102. doi:10.1016/j.devcel.2014.06.007

77. Hönscher C, Mari M, Auffarth K, Bohnert M, Griffith J, Geerts W, et al. Short Article. Dev Cell. Elsevier Inc; 2014;30: 86–94. doi:10.1016/j.devcel.2014.06.006

78. Bond S, Forgac M. The Ras/cAMP/protein kinase A pathway regulates glucose-dependent assembly of the vacuolar (H+)-ATPase in yeast. J Biol Chem. American Society for Biochemistry and Molecular Biology; 2008;283: 36513–36521. doi:10.1074/jbc.M805232200

79. Nelson H, Nelson N. Disruption of genes encoding subunits of yeast vacuolar H(+)-ATPase causes conditional lethality. Proc Natl Acad Sci USA. 1990;87: 3503–3507. doi:10.2307/2354196

80. Yamashiro CT, Kane PM, Wolczyk DF, Preston RA, Stevens TH. Role of vacuolar acidification in protein sorting and zymogen activation: a genetic analysis of the yeast vacuolar proton-translocating ATPase. Mol Cell Biol. American Society for Microbiology (ASM); 1990;10: 3737–3749.

81. Thorsness PE, White KH, Fox TD. Inactivation of YME1, a member of the ftsH-SEC18-PAS1-CDC48 family of putative ATPase-encoding genes, causes increased escape of DNA from mitochondria in Saccharomyces cerevisiae. Mol Cell Biol. 1993;13: 5418–5426.

82. Dunn CD, Tamura Y, Sesaki H, Jensen RE. Mgr3p and Mgr1p are adaptors for the mitochondrial i-AAA protease complex. Mol Biol Cell. 2008 ed. 2008;19: 5387–5397. doi:10.1091 /mbc. E08-01-0103

83. Ieva R, Schrempp SG, Opaliński L, Wollweber F, Höß P, Heißwolf AK, et al. Mgr2 functions as lateral gatekeeper for preprotein sorting in the mitochondrial inner membrane. Mol Cell. 2014;56: 641–652. doi:10.1016/j.molcel.2014.10.010

84. Artal-Sanz M, Tavernarakis N. Prohibitin and mitochondrial biology. Trends in Endocrinology & Metabolism. 2009;20: 394–401. doi:10.1016/j.tem.2009.04.004

85. Kominsky DJ, Brownson MP, Updike DL, Thorsness PE. Genetic and biochemical basis for viability of yeast lacking mitochondrial genomes. Genetics. 2002;162: 1595–1604.

86. Appleby RD, Porteous WK, Hughes G, James AM, Shannon D, Wei YH, et al. Quantitation and origin of the mitochondrial membrane potential in human cells lacking mitochondrial DNA. Eur J Biochem. 1999;262: 108–116.

87. Chen WW, Birsoy K, Mihaylova MM, Snitkin H, Stasinski I, Yucel B, et al. Inhibition of ATPIF1 ameliorates severe mitochondrial respiratory chain dysfunction in mammalian cells. CellReports. 2014;7: 27–34. doi:10.1016/j.celrep.2014.02.046

88. Chen XJ, Hansbro PM, Clark-Walker GD. Suppression of rho0 lethality by mitochondrial ATP synthase F1 mutations in Kluyveromyces lactis occurs in the absence of F0. Mol Gen Genet. 1998;259: 457–467.

89. Kovacova V, Irmlerova J, Kovac L. Oxidative phosphorylatiion in yeast. IV. Combination of a nuclear mutation affecting oxidative phosphorylation with cytoplasmic mutation to respiratory deficiency. Biochim Biophys Acta. 1968;162: 157–163.

90. Chen XJ. Sal1p, a calcium-dependent carrier protein that suppresses an essential cellular function associated With the Aac2 isoform of ADP/ATP translocase in *Saccharomyces cerevisiae*. Genetics. 2004;167: 607–617. doi:10.1534/genetics. 103.023655

91. Pfanner N, Neupert W. Transport of F1-ATPase subunit beta into mitochondria depends on both a membrane potential and nucleoside triphosphates. FEBS Lett. 1986;209: 152–156.

92. Kang PJ, Ostermann J, Shilling J, Neupert W, Craig EA, Pfanner N. Requirement for hsp70 in the mitochondrial matrix for translocation and folding of precursor proteins. Nature. Nature Publishing Group; 1990;348: 137–143. doi:10.1038/348137a0

93. Dunn CD, Jensen RE. Suppression of a defect in mitochondrial protein import identifies cytosolic proteins required for viability of yeast cells lacking mitochondrial DNA. Genetics. 2003rd ed. 2003;165: 35–45.

94. Kerscher O, Sepuri NB, Jensen RE. Tim18p is a new component of the Tim54p-Tim22p translocon in the mitochondrial inner membrane. Mol Biol Cell. 2000 ed. 2000;11: 103–116.

95. Rahman MU, Hudson AP. Substrates for yeast mitochondrial cAMP-dependent protein kinase activity. Biochemical and Biophysical Research Communications. 1995;214: 188–194. doi:10.1006/bbrc.1995.2273

96. Gerbeth C, Schmidt O, Rao S, Harbauer AB, Mikropoulou D, Opalinska M, et al. Glucose-induced regulation of protein import receptor Tom22 by cytosolic and mitochondria-bound kinases. Cell metabolism. 2013;18: 578–587. doi:10.1016/j.cmet.2013.09.006

97. Rao S, Schmidt O, Harbauer AB, Schönfisch B, Guiard B, Pfanner N, et al. Biogenesis of the preprotein translocase of the outer mitochondrial membrane: protein kinase A phosphorylates the precursor of Tom40 and impairs its import. Mol Biol Cell. American Society for Cell Biology; 2012;23: 1618–1627. doi:10.1091/mbc.E11-11-0933

98. Schmidt O, Harbauer AB, Rao S, Eyrich B, Zahedi RP, Stojanovski D, et al. Regulation of mitochondrial protein import by cytosolic kinases. Cell. 2011; 144: 227–239. doi:10.1016/j.cell.2010.12.015

99. Ryan MT, Müller H, Pfanner N. Functional staging of ADP/ATP carrier translocation across the outer mitochondrial membrane. J Biol Chem. 1999;274: 20619–20627.

100. Wiedemann N, Pfanner N, Ryan MT. The three modules of ADP/ATP carrier cooperate in receptor recruitment and translocation into mitochondria. EMBO J. EMBO Press; 2001; 20: 951–960. doi:10.1093/emboj/20.5.951

101. Dietmeier K, Zara V, Palmisano A, Palmieri F, Voos W, Schlossmann J, et al. Targeting and translocation of the phosphate carrier/p32 to the inner membrane of yeast mitochondria. J Biol Chem. 1993;268: 25958–25964.

102. Dirick L, Bendris W, Loubiere V, Gostan T, Gueydon E, Schwob E. Metabolic and environmental conditions determine nuclear genomic instability in budding yeast lacking mitochondrial DNA. G3 (Bethesda). Genetics Society of America; 2014;4: 411–423. doi:10.1534/g3.113.010108

103. Birsoy K, Possemato R, Lorbeer FK, Bayraktar EC, Thiru P, Yucel B, et al. Metabolic determinants of cancer cell sensitivity to glucose limitation and biguanides. Nature. Nature Publishing Group; 2014;508: 108–112. doi:10.1038/nature13110

104. Chevtzoff C, Vallortigara J, Avéret N, Rigoulet M, Devin A. The yeast cAMP protein kinase Tpk3p is involved in the regulation of mitochondrial enzymatic content during growth. Biochim Biophys Acta. 2005;1706: 117–125. doi:10.1016/j.bbabio.2004.10.001

105. Leadsham JE, Gourlay CW. cAMP/PKA signaling balances respiratory activity with mitochondria dependent apoptosis via transcriptional regulation. BMC Cell Biol. BioMed Central Ltd; 2010;11: 92. doi:10.1186/1471-2121-11-92

106. Demlow CM, Fox TD. Activity of mitochondrially synthesized reporter proteins is lower than that of imported proteins and is increased by lowering cAMP in glucose-grown Saccharomyces cerevisiae cells. Genetics. 2003;165: 961–974.

107. Bodenmiller B, Wanka S, Kraft C, Urban J, Campbell D, Pedrioli PG, et al. Phosphoproteomic Analysis Reveals Interconnected System-Wide Responses to Perturbations of Kinases and Phosphatases in Yeast. Science signaling. 2010;3: rs4–rs4. doi:10.1126/scisignal.2001182

108. Brett CL, Kallay L, Hua Z, Green R, Chyou A, Zhang Y, et al. Genome-Wide Analysis Reveals the Vacuolar pH-Stat of *Saccharomyces cerevisiae*. Kaeberlein M, editor. PLoS ONE. 2011; 6: e17619. doi:10.1371/journal.pone.0017619.g004

109. Dimitrov LN, Brem RB, Kruglyak L, Gottschling DE. Polymorphisms in multiple genes contribute to the spontaneous mitochondrial genome instability of *Saccharomyces cerevisiae* S288C strains. Genetics. 2009;183: 365–383. doi:10.1534/genetics. 109.104497

110. Yang Y, Foulquié-Moreno MR, Clement L, Erdei E, Tanghe A, Schaerlaekens K, et al. QTL analysis of high thermotolerance with superior and downgraded parental yeast strains reveals new minor QTLs and converges on novel causative alleles involved in RNA processing. Steinmetz LM, editor. PLoS Genet. Public Library of Science; 2013;9: e1003693. doi:10.1371/journal.pgen.1003693

111. Gaisne M, Bécam AM, Verdière J, Herbert CJ. A “natural” mutation in Saccharomyces cerevisiae strains derived from S288c affects the complex regulatory gene HAP1 (CYP1). Curr Genet. 1999;36: 195–200.

112. Birsoy K, Wang T, Chen WW, Freinkman E, Abu-Remaileh M, Sabatini DM. An Essential Role of the Mitochondrial Electron Transport Chain in Cell Proliferation Is to Enable Aspartate Synthesis. Cell. 2015;162: 540–551. doi:10.1016/j.cell.2015.07.016

113. Sullivan LB, Gui DY, Hosios AM, Bush LN, Freinkman E, Vander Heiden MG. Supporting Aspartate Biosynthesis Is an Essential Function of Respiration in Proliferating Cells. Cell. 2015;162: 552–563. doi:10.1016/j.cell.2015.07.017

114. Koga Y, Povalko N, Nishioka J, Katayama K, Yatsuga S, Matsuishi T. Molecular pathology of MELAS and L-arginine effects. Biochim Biophys Acta. 2012;1820: 608–614. doi:10.1016/j.bbagen.2011.09.005

115. Jauniaux JC, Urrestarazu LA, Wiame JM. Arginine metabolism in *Saccharomyces cerevisiae:* subcellular localization of the enzymes. J Bacteriol. American Society for Microbiology (ASM); 1978;133: 1096–1107.

116. Oldenburg KR, Vo KT, Michaelis S, Paddon C. Recombination-mediated PCR-directed plasmid construction in vivo in yeast. Nucleic Acids Res. 1997;25: 451–452.

117. Afgan E, Baker D, Coraor N, Goto H, Paul IM, Makova KD, et al. Harnessing cloud computing with Galaxy Cloud. Nat Biotechnol. Nature Publishing Group; 2011;29: 972–974. doi:10.1038/nbt.2028

118. Trapnell C, Roberts A, Goff L, Pertea G, Kim D, Kelley DR, et al. Differential gene and transcript expression analysis of RNA-seq experiments with TopHat and Cufflinks. Nat Protoc. Nature Publishing Group; 2012;7: 562–578. doi:10.1038/nprot.2012.016

119. Trapnell C, Hendrickson DG, Sauvageau M, Goff L, Rinn JL, Pachter L. Differential analysis of gene regulation at transcript resolution with RNA-seq. Nat Biotechnol. 2013;31: 46–53. doi:10.1038/nbt.2450

120. Engel SR, Dietrich FS, Fisk DG, Binkley G, Balakrishnan R, Costanzo MC, et al. The Reference Genome Sequence of *Saccharomyces cerevisiae:* Then and Now. G3 (Bethesda). 2014;4: 389–398. doi:10.1534/g3.113.008995

121. Cherry JM, Hong EL, Amundsen C, Balakrishnan R, Binkley G, Chan ET, et al. Saccharomyces Genome Database: the genomics resource of budding yeast. Nucleic Acids Res. Oxford University Press; 2012;40: D700–5. doi:10.1093/nar/gkr1029

122. Orlova M, Barrett L, Kuchin S. Detection of endogenous Snf1 and its activation state: application to Saccharomycesand Candidaspecies. Yeast. 2008;25: 745–754. doi:10.1002/yea.1628

123. Updyke TV, Engelhorn SC, Novel Experimental Technology, Inc. System for pH-neutral stable electrophoresis gel. US Patent Office; US 6162338 A, 1999.

124. Rutherford JC. Aft1p and Aft2p Mediate Iron-responsive Gene Expression in Yeast through Related Promoter Elements. 2003;278: 27636–27643. doi:10.1074/jbc. M300076200

125. Puig S, Askeland E, Thiele DJ. Coordinated Remodeling of Cellular Metabolism during Iron Deficiency through Targeted mRNA Degradation. Cell. 2005;120: 99–110. doi:10.1016/j.cell.2004.11.032

126. DeRisi J, van den Hazel B, Marc P, Balzi E, Brown P, Jacq C, et al. Genome microarray analysis of transcriptional activation in multidrug resistance yeast mutants. FEBS Lett. Elsevier Science; 2000;470: 156–160.

127. Devaux F, Marc P, Bouchoux C, Delaveau T, Hikkel I, Potier MC, et al. An artificial transcription activator mimics the genome-wide properties of the yeast Pdr1 transcription factor. EMBO Rep. 2001; 2: 493–498. doi:10.1093/embo-reports/kve114

128. Pavlidis P, Noble WS. Matrix2png: a utility for visualizing matrix data. Bioinformatics. 2003;19: 295–296.

129. Sikorski RS, Hieter P. A system of shuttle vectors and yeast host strains designed for efficient manipulation of DNA in *Saccharomyces cerevisiae*. Genetics. Genetics Society of America; 1989;122: 19–27.

130. Sesaki H, Jensen RE. Division versus fusion: Dnm1p and Fzo1p antagonistically regulate mitochondrial shape. J Cell Biol. 1999;147: 699–706.

131. Breker M, Gymrek M, Schuldiner M. A novel single-cell screening platform reveals proteome plasticity during yeast stress responses. J Cell Biol. Rockefeller Univ Press; 2013;200: 839–850. doi:10.1083/jcb.201301120

132. Bähler J, Wu JQ, Longtine MS, Shah NG, McKenzie A, Steever AB, et al. Heterologous modules for efficient and versatile PCR-based gene targeting in Schizosaccharomyces pombe. Yeast. John Wiley & Sons, Ltd; 1998;14: 943–951. doi:10.1002/(SICI)1097-0061 (199807)14:10<943::AID-YEA292>3.0.CO;2-Y

133. Taxis C, Knop M. System of centromeric, episomal, and integrative vectors based on drug resistance markers for Saccharomyces cerevisiae. BioTechniques. 2006 Jan pp. 73–78.

